# Toward pathology-guided illumination optimization for neuromodulation with a foundation-model-encoded light transport surrogate

**DOI:** 10.64898/2026.07.04.736179

**Authors:** Sihan Dong, Maoye Guan, Liqin Yang, Guoquan Liu, Axel Rominger, Yixuan Yuan, Wuwei Ren, Ruiqing Ni, Xunbin Wei

**Author notes:** Correspondence to: Xunbin Wei,; Ruiqing Ni,; Wuwei Ren.

## Abstract

Clinical treatment planning for near-infrared (NIR) light neuromodulation requires patient-specific dosimetry to optimize light fluence (LF) delivery to cortical targets. The gold-standard Monte Carlo photon-transport forward solver is accurate but computationally expensive and non-differentiable for personalized inverse design across subjects. Here, we present PhomiNeuro, a foundation-model-encoded differentiable surrogate for time-resolved LF modeling and pathology-guided inverse design. A pretrained 3D medical imaging foundation model (VISTA3D) was domain-adapted to 285 training head models annotated with optical properties, and then coupled to an implicit neural representation (INR) that predicts LF at arbitrary spatiotemporal coordinates. Regularized by a time-dependent diffusion-equation residual, it demonstrated superior fidelity in 84 held-out participants, and ablations identified VISTA3D-derived anatomical priors as dominant and physics regularization as complementary. PhomiNeuro enables fast and robust queries of LF and its gradients with respect to illumination parameters for targets guided by individual structural magnetic resonance imaging and amyloid positron emission tomography, achieving 466-fold per-iteration speedup and 167-fold end-to-end optimization speedup. Explanatory analyses of LF delivery variability, including sex differences and amyloid status, identified structural variations as the primary drivers. These results position PhomiNeuro as a highly extensible translational framework toward personalized treatment planning and digital twin development in precision neuromodulation.

## Introduction

Near-infrared (NIR) light brain stimulation, also known as transcranial photobiomodulation (tPBM), is a promising non-invasive intervention for neurodegenerative disorders, including Alzheimer’s disease (AD)^1,2^ and Parkinson’s disease^3^. In preclinical AD models, tPBM has been shown to reduce cerebral amyloid-β (Aβ) burden, attenuate pro-inflammatory microglial responses, promote angiogenesis, and enhance meningeal lymphatic drainage, with associated improvements in cognitive function^4–6^. Emerging clinical evidence further supports its cognitive benefits, favorable safety profile, high treatment adherence, and the feasibility of home-based administration^7–9^. Realizing this therapeutic potential requires accurate intracranial light dosimetry and adaptive treatment planning^10^. Currently, its clinical translation remains hindered by fixed protocols that lack anatomy- and pathology-informed personalization^11^. As light is strongly scattered and absorbed when propagating through scalp, skull, cerebral vasculature, and brain parenchyma, only a small fraction of incident energy reaches intracranial targets^12^. Interindividual anatomical variability, including sex- and AD-associated differences in scalp and skull thickness, cerebrospinal fluid (CSF) anatomy, and brain parenchymal fraction (BPF), further alters transcranial light delivery^13^. These challenges motivate personalized optimization of source placement, orientation, and beam parameters to maximize light fluence (LF) within therapeutic targets while satisfying safety constraints^14^.

Recent efforts toward personalized tPBM includes anatomy-based light transport modeling^15^, noninvasive optical measurements of target-region LF^16^, and treatment-response monitoring using functional near-infrared spectroscopy (fNIRS)^17,18^. Inspired by radiotherapy where Monte Carlo (MC) simulation is a gold standard for patient-specific dosimetry and precision treatment planning^19^, tPBM has increasingly adopted MC methods as reference models for accurate photon-transport modeling in heterogeneous head tissues^20^. However, it still lacks theranostic frameworks linking imaging-defined targets and patient-specific dosimetry to optimized treatment planning analogous to those in radiotherapy^21^ and nuclear medicine^22^. As evaluating candidate illumination settings requires repeated simulations, while reliable estimates in weakly illuminated regions require substantial photon sampling, personalized illumination planning is computationally prohibitive. Despite that parallel computing accelerates MC methods^23,24^, it does not eliminate the trade-off between computation time and statistical precision. Although learning-based surrogates^25,26^ and denoisers^27^ offer accelerated or enhanced predictions with reduced noise, current methodologies primarily predict light distributions within simplified geometric models or standard head atlases, which severely limits their generalizability. Moreover, these methods only operate under a specified source configuration, failing to explicitly indicate how illumination parameters should be altered to improve target LF delivery. Therefore, personalized tPBM planning requires a strategy that preserves MC fidelity, enables generalization across individual anatomies, and provides differentiable inference with sensitivities to illumination parameters.

Here, we present PhomiNeuro (**Fig. 1a**), an open-source framework that combines the complementary strengths of a medical imaging foundation model (FM) and an implicit neural representation (INR) to enable time-resolved light transport modeling and pathology-guided illumination planning. The FM provides the anatomical backbone by leveraging multiscale priors learned from large, anatomically diverse volumetric datasets to support generalization across participants, acquisition protocols, and downstream tasks with limited task-specific supervision^28–31^. In contrast, the INR is well suited as a light transport solver that represents continuous, source-conditioned LF fields at arbitrary coordinates, whereas conventional fixed-grid models often blur sharp near-source field peaks^32,33^. By parameterizing LF as a function of spatiotemporal coordinates and illumination parameters, the INR enables differentiation of target LF with respect to source settings at arbitrary coordinates. The time-resolved architecture can also extend dosimetry beyond steady-state exposure to emerging pulsed tPBM paradigms, for which stimulation frequency and pulse duration can elicit distinct biological responses^4,34^.

**Fig. 1:**
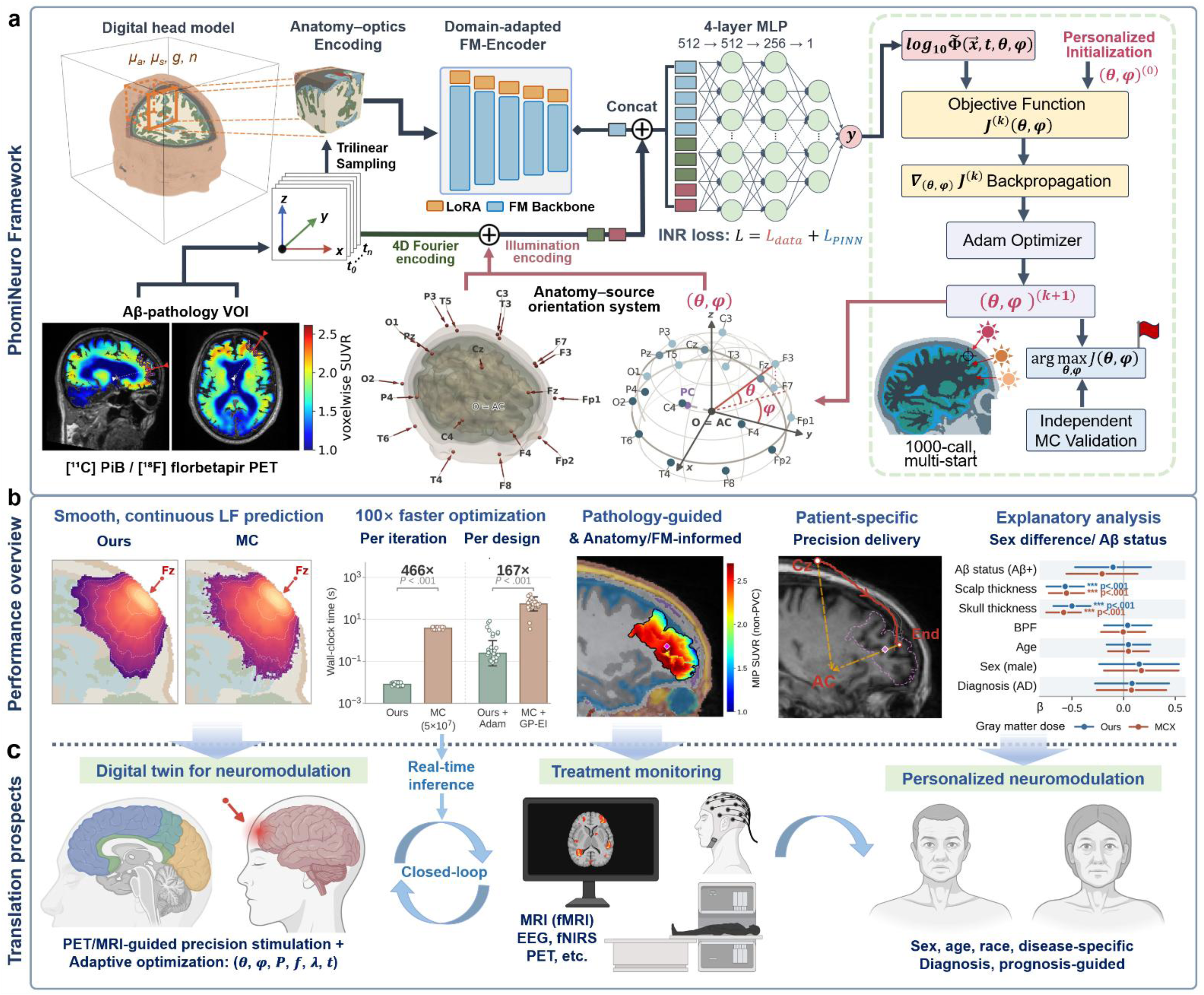
Overview of our PhomiNeuro framework for personalized illumination optimization and pathology-guided neuromodulation planning. (a) Forward modeling and inverse optimization of light fluence (LF) field with PhomiNeuro. After adaption to anatomical and tissue-optics domain of the training head models by masked autoencoding with low-rank adaptation (LoRA, rank 32, *α* = 64), the encoder of a pretrained 3D medical foundation model (FM) is used to extract local anatomical-optical features via trilinear sampling at a query coordinate (*x*, *y*, *z*). Then an implicit neural representation (INR) model is trained to predict the LF log_10_ Φ^(*x*, *y*, *z*, *t*) at time gate *t*. Its input comprises spatiotemporal coordinate features and illumination parameters, which concatenates FM-encoded five-scale features, band-limited Fourier encodings of (*x*, *y*, *z*, *t*), and light beam direction. The INR loss comprises a physics-informed term *L_PINN_* and three data terms, where the *L_PINN_* is the normalized residual of the time-dependent diffusion equation. Individualized pathology-guided optimization of tPBM was performed guided by amyloid-PET SUVR; an automatic gradient-based, multi-start Adam optimizer is designed for the inverse optimization. See Fig S11 for details. (**b**) Performance overview. PhomiNeuro enables accurate, off-grid LF prediction, with suppressed discrete noise and 466-fold speedup per iteration and 167-fold speedup for the entire inverse optimization process relative to the Monte Carlo (MC) method. The inverse design can be guided by amyloid-PET pathology and FM-encoded anatomy information for precision energy delivery. Explanatory analysis for inter-subject variability, including sex difference and amyloid-status was performed both on our model and MC. (**c**) Clinical translation Perspectives. ***θ***, ***φ***, source orientation; ***P***, ***f***, ***λ***, ***t***, stimulation parameters: power density, repetition frequency, wavelength, duration. Abbreviations: Aβ, Amyloid-beta; AC, Anterior Commissure; Adam, Adaptive Moment Estimation; AD, Alzheimer’s disease; BPF, Brain Parenchyma Fraction; EEG, Electroencephalography; fMRI, Functional Magnetic Resonance Imaging; fNIRS, Functional Near-Infrared Spectroscopy; MLP, Multilayer Perceptron; PET, Positron Emission Tomography; SUVR, Standardized Uptake Value Ratio; VOI, Volume of Interest. The 3D phantoms shown in (a) are derived from a publicly distributed BrainWeb digital head model and the MNI-ICBM152 average template. Created with Python, Microsoft PowerPoint and BioRender.com.

Specifically, PhomiNeuro adapts the pretrained VISTA3D FM^35^ to the optical-property domain through parameter-efficient fine-tuning with a masked-autoencoding objective^36,37^ on 285 training head models, thereby enabling its learned multiscale anatomical representations to encode participant-specific distributions of tissue optical properties. At each query coordinate, local features sampled from the resulting multiscale FM representation are coupled to a lightweight, source-conditioned INR, enabling fast LF and gradient queries for differentiable optimization of illumination parameters for targets defined by positron emission tomography (PET). This light transport surrogate predicts the time-resolved LF response to an impulsive source using 0.2-ns time gating, allowing the derivation of temporal point spread function (tPSF). In addition, a band-limited Fourier encoding^38^ is leveraged to preserve rapid near-source variation and time-gated temporal dynamics, while a residual of the time-dependent diffusion equation (TD DE) is employed to promotes physical consistency and suppresses MC sampling noise. This factorization separates participant-specific anatomy from source- and coordinate-dependent light transport, allowing the multiscale anatomical representation to be computed once per head and reused for fast forward dosimetry and continuous inverse design.

Across 84 held-out participants, PhomiNeuro accurately predicted time-resolved LF and reproduced the structural determinants of interparticipant dose variability identified by MC (**Fig. 1b**). Its FM-based anatomical encoding, continuous field representation, and physics-informed regularization collectively improved performance over the benchmark models. As an MC surrogate, PhomiNeuro further enabled fast, noise-suppressed, off-grid LF inference and accelerated light-source orientation optimization by two orders of magnitude to achieve sub-second computation times across 28 amyloid-positive participants. The framework used amyloid PET to define tPBM targets from the spatial distribution of hallmark Aβ pathology and structural magnetic resonance imaging (MRI) to model participant-specific whole-head anatomy, including AD-related brain atrophy and variations in scalp thickness. Explanatory analyses further identified structural, pathological, and sex-related factors associated with LF distribution patterns, with these associations corroborated by the MC simulations. More broadly, PhomiNeuro may provide a computationally efficient foundation for a future tPBM digital twin^39^ (**Fig. 1c**) that combines multimodal imaging-derived whole-head anatomy, brain function, and pathology with patient-specific optical dosimetry, differentiable optimization, and longitudinal response monitoring. Such a digital twin could dynamically update pathology-defined targets and illumination parameters using patient-specific imaging and physiological feedback, thereby linking diagnosis, treatment planning, intervention delivery, and response assessment into a closed-loop optical theranostic workflow for personalized neuromodulation across diverse neurodegenerative disorders.

## Results

### PhomiNeuro achieves accurate and robust time-resolved light transport prediction

PhomiNeuro accurately captured the spatiotemporal evolution of the diffuse fluence distribution, as visually evaluated using a development-test head model under frontal-midline illumination (**Fig. 2a**, **Video S1**). The predicted field also preserved the LF propagating pattern along CSF lateral to the beam direction, consistent with the MC-resolved diffusive waveguide effect which enhances light transmission through the CSF surrounding the brain^40^. In another head model under right-temporal illumination (**Fig. 2b**), the cross-sectional evaluation demonstrated suppressed discrete speckle noise in the CSF comparing to MC. Visual evaluation of our time-resolved LF prediction and prediction residuals against MC were benchmarked versus five grid-based baseline networks, including VIT^41^, DynUNet^42^, SegResNet^43^, UNet^44^, a Fourier neural operator (FNO)^45^, and two coordinate baseline networks: a random-Fourier-feature (RFF) multilayer perceptron^38^ and SIREN^46^ (**Fig. 2c**). The LF field predicted by PhomiNeuro provided the highest fidelity to MC reference LF field, showing lower energy leakage and higher smoothness in dark region of deep tissue versus the coordinate networks, and lower peak LF residual across the time range 0-1.2 ns comparing to the grid-based networks. In addition to the 20 development-test normal participants involved in architecture selection and ablation, final evaluation was benchmarked using 84 held-out OASIS-3 participants, including 42 participants with AD and 42 marched cognitively unimpaired (CU) participants. Across the 84 held-out participants, PhomiNeuro achieved the highest gate-averaged R² (0.7385 ± 0.0019) and lowest Root Mean Squared Error (RMSE, 0.6211 ± 0.0027 decades) among the eight networks (**Fig. 2d**, **Table S1**). It also preserved LF field contrast more faithfully with a variance ratio of 0.819, and achieved the highest Lin concordance correlation coefficient of 0.863 (**Fig. 2e**). Gate-specific R² and RMSE analyses in the held-out cohort showed that PhomiNeuro outperformed all baselines during 0-1.0 ns (**Tables S2**, **S3**), which is the principal energy-delivery window accounting for approximately 85% of the time-integrated whole-brain absorbed energy (**Fig. S3**).

**Fig. 2:**
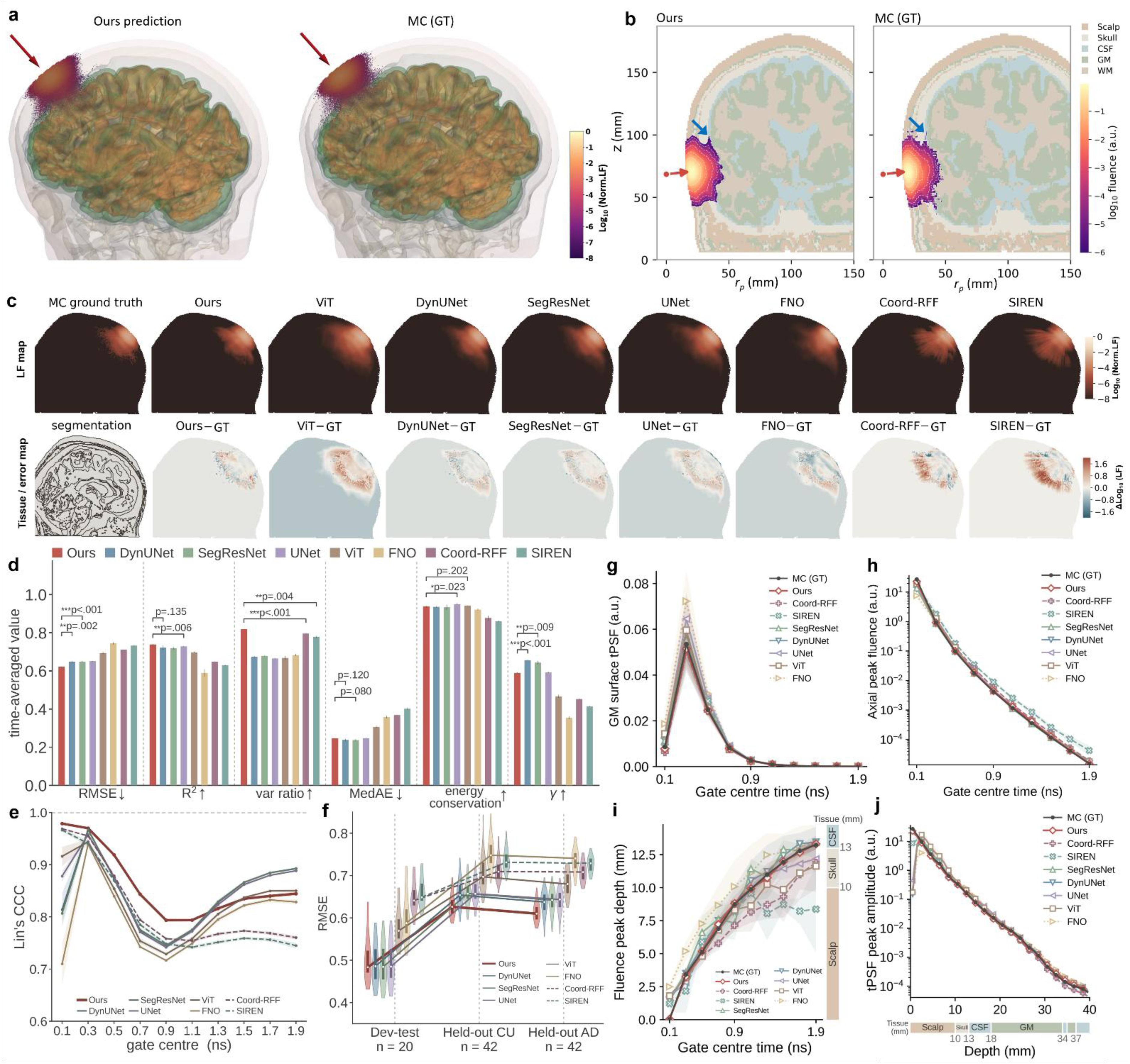
Benchmark of our light transport surrogate performance against other baseline networks. **(a)** 3D rendering of a development-test (dev-test) head model (frontal midline, Fz; time gate, 0.2–0.4 ns) from the scatterBrains dataset. Light fluence (LF) fields from our prediction and Monte Carlo (MC) ground truth are shown with particle density. Red arrow, beam direction; colormap, normalized log_10_*Φ*. (**b**) LF within a cross-section through the beam direction for a dev-test head model (right temporal, T4; time gate 0.2–0.4 ns) from the BrainWeb dataset. Blue arrows indicate discrete noise suppressed by PhomiNeuro in the cerebrospinal fluid (CSF) region compared to the MC method. (**c**) LF fields and LF prediction residuals on a dev-test head model at time gate 0.4–0.6 ns. First row: LF in the plane through the beam, MC ground truth, ours and seven baseline networks. Second row: tissue segmentation and LF residual of each network against the MC ground truth. (**d**) Benchmark on the held-out cohort (n=84, OASIS head models, 6 stimulation sites, three seeds per network, averaged over time gates) across six metrics: RMSE, *R*^2^, variance ratio, energy deposition conservation, median absolute error (MedAE), and γ (3%/2 px) pass rate. Error bars, s.d. across training seeds (3 seeds per model). (**e**) Lin’s concordance correlation coefficient (CCC) per time gate. Shading, ±1 s.e. calculated by combining a 2,000-fold bootstrap over the 84 subjects with the between-seed s.d. (**f**) Generalization across cohorts. RMSE distributions for head models of participants from the dev-test cohort (n=20), held-out cognitive unimpaired (CU) cohort (n=42) and held-out AD cohort (n=42). (**g-j**) Temporal and spatial distribution of LF prediction and MC GT on the beam direction axis at the T3 stimulation site on the held-out cohort (n=84, three seeds per model): (**g**) temporal point-spread function (tPSF) at the grey-matter surface voxel on the beam axis; (**h**) amplitude of the LF maximum versus gate time; (**i**) penetration depth of the LF peak versus time; (**j**) tPSF peak amplitude versus penetration depth below scalp surface. Curves, mean over the 84 subjects; shading, ±1 s.e. calculated by combining the standard error across participants and the between-seed s.d.

Cross-cohort evaluation further distinguished the generalization of these networks (**Fig. 2f, Table S4**). Although PhomiNeuro did not outperform SegResNet, DynUNet, and UNet on the development-test set, it maintained robust performance to lead all models in the unseen held-out cohort, whereas these baselines exhibited degradation much more markedly. Coord-RFF transferred without deterioration but remained substantially less accurate. All networks showed slightly lower RMSE in AD than in CU participants, including ours (0.610 vs. 0.626), suggesting that this small difference reflects cohort anatomy rather than disease-specific model behavior. PhomiNeuro also accurately reproduced the spatiotemporal dynamics of the diffuse LF field across the 84 held-out participants (**Fig. 2g-j**): the peak LF amplitude and depth were recovered with mean absolute errors of 0.027 decades and 0.114 mm respectively at the left temporal region; the peak-depth error was 0.020 mm at the first gate (0-0.2 ns); the peak-amplitude error remained 0.073 decades at the first voxel beneath the scalp, where the LF gradient was steepest; and the time-to-peak error remained below 0.086 ns throughout the evaluated depth range.

The continuous representation was further evaluated beyond the native temporal and angular sampling grids (**Fig. S4**). At intermediate time points absent from training, PhomiNeuro recovered the non-linear tPSF obtained from an independent 40-gate MC simulation and generated smooth LF distributions at interpolation time point such as 0.35, 0.40, and 0.45 ns (**Video S2**). Along an 80-mm depth profile, the time-integrated LF prediction remained within −0.33 to +0.64 decades of the higher sampling MC reference with 5×10^9^ photons, whereas the standard 5×10⁷-photon MC simulation increasingly underestimated the deep LF as photon counts diminished. When extrapolating beyond the trained beam-elevation range, PhomiNeuro maintained RMSE values of 0.335–0.436 decades above the upper boundary (R² = 0.872−0.924), whereas performance below the lower boundary varied more strongly with azimuth (RMSE = 0.253−1.036 decades; R² = 0.505−0.957). The standard MC simulation remained more accurate in the seven brightest LF bands, while no significant difference was detected in the two darkest bands (paired Wilcoxon P = 0.13 and 0.25). Thus, the principal advantage of PhomiNeuro in this regime lies in its smooth, continuous, and off-grid query capability rather than superior fidelity to MC under matched reference conditions.

### Ablation studies identify FM encoding as dominant and physics regularization as complementary

The pretrained and domain-adapted FM-encoder provides anatomical-optical conditioning that is identified as the principal contributor to predictive accuracy by prespecified ablation on the development-test cohort (**Table 1**). Removing the adapted FM encoder reduced R² from 0.7632 ± 0.0039 to 0.6508 ± 0.0047 and increased RMSE from 0.6491 ± 0.0050 to 0.7807 ± 0.0043. Replacing it with a randomly initialized encoder produced a similar loss (R² = 0.6785 ± 0.0069), whereas retaining the pretrained encoder without domain adaptation caused a modest decrease (R² = 0.7557 ± 0.0008), indicating that the pretrained anatomical features provided the major improvement and optical-property adaptation refined their alignment with photon transport in head tissues. Removing source-to-query path integrals produced a smaller but consistent reduction in R² to 0.7416 ± 0.0067 and in gamma pass rate from 0.5835 ± 0.0040 to 0.5202 ± 0.0045.

**Table 1:** Ablation experiment of PhomiNeuro on the development cohort (20 dev-test head models with 19 stimulation sites, three seeds per variant).

| **Variant** | **R²** | **RMSE** | **MedAE** | **Spearman ρ** | **γ (3%/2 px)** |
|---|---|---|---|---|---|
| **PhomiNeuro** | **0.7632 ± 0.0039** | **0.6491 ± 0.0050** | 0.2844 ± 0.0008 | **0.9250 ± 0.0004** | 0.5835 ± 0.0040 |
| − FM encoder | 0.6508 ± 0.0047 | 0.7807 ± 0.0043 | 0.3620 ± 0.0027 | 0.8915 ± 0.0008 | 0.4429 ± 0.0032 |
| FM encoder (not fine-tuned) | 0.7557 ± 0.0008 | 0.6594 ± 0.0022 | 0.2942 ± 0.0006 | 0.9231 ± 0.0005 | 0.5560 ± 0.0025 |
| Random encoder | 0.6785 ± 0.0069 | 0.7516 ± 0.0070 | 0.3605 ± 0.0016 | 0.9054 ± 0.0007 | 0.5131 ± 0.0047 |
| − path integrals | 0.7416 ± 0.0067 | 0.6796 ± 0.0078 | 0.3184 ± 0.0003 | 0.9182 ± 0.0009 | 0.5202 ± 0.0045 |
| − loss term *L<sub>PINN</sub>* | 0.6874 ± 0.0009 | 0.7333 ± 0.0008 | 0.2946 ± 0.0006 | 0.9174 ± 0.0021 | 0.5993 ± 0.0020 |
| − loss term *L<sub>lin</sub>* | 0.7612 ± 0.0007 | 0.6528 ± 0.0006 | 0.2836 ± 0.0011 | 0.9248 ± 0.0007 | 0.5758 ± 0.0021 |
| − loss term *L<sub>hinge</sub>* | 0.6103 ± 0.0631 | 0.8459 ± 0.0570 | 0.4474 ± 0.0127 | 0.8731 ± 0.0045 | 0.2623 ± 0.0831 |
| −*L<sub>PINN</sub>* and − FM encoder | 0.6185 ± 0.0025 | 0.8084 ± 0.0017 | 0.3662 ± 0.0012 | 0.8870 ± 0.0010 | 0.4492 ± 0.0020 |

Mean ± across-seed s.d. over three training seeds, for the complete model and for every ablated arm alike; the s.d. is the spread of the gate-averaged value across seeds. A difference from the complete model is outside the noise when it exceeds $1.96\sqrt{((\sigma_h{}^2 + \sigma_v{}^2)/3)}$. A same posteriori ablation on the held-out cohort can be found in Table S5.

Ablation study of the individual loss terms revealed a clearer hierarchy: Removing the censored hinge loss, which constrains predictions in photon-starved deep-tissue regions with sparse MC sampling, caused the largest degradation in development-set performance (R² = 0.6103 ± 0.0631; RMSE = 0.8459 ± 0.0570); while removing the linear-space term had little effect on aggregate accuracy (R² = 0.7612 ± 0.0007). Excluding the physic-informed TD DE residual reduced development R² to 0.6874 ± 0.0009. Joint removal of the FM encoder and physics residual yielded the lowest accuracy among these representation ablations (R² = 0.6185 ± 0.0025), supporting complementary contributions from multiscale anatomy and physics-informed regularization during model development. Post-hoc analysis in the held-out cohort further confirmed the importance of FM-encoded anatomical-optical conditioning, path integrals, and censored supervision (**Table S5**). Removing the FM encoder or path integrals reduced predictive accuracy, while eliminating the hinge loss caused the largest R² collapse. By contrast, removing the physics residual or linear-space term did not reduce aggregate held-out accuracy. A separate temporal-encoding ablation supported the final three-octave, Nyquist-consistent design: native-gate accuracy remained unchanged, whereas intermediate-gate R² increased from 0.6044 to 0.7413.

### Differentiable inverse design enables fast pathology-guided illumination optimization

Pathology-guided targets were defined from participant-specific amyloid PET within DKT-atlas-based cortical parcellations (**Fig. 3a-b**). The analysis included 28 held-out participants with session-matched PET and T1-weighted (T1w) MRI, comprising 14 AD and 14 CU participants with Centiloid values above 26 (defined as amyloid-positive, Aβ+)^47,48^. Voxelwise standardized uptake value ratios (SUVR) were projected onto each participant’s cortical surface, while partial-volume-corrected regional values were mapped onto the parcellated T1w volume. The Aβ burden-weighted centroid of the highest-burden cortical parcel within a 32-mm radial depth defined the targets, which occurred most frequently in the rostral middle frontal gyrus, banks of the superior temporal sulcus, and frontal pole, at depths of 18.3–31.5 mm (**Fig. S5, S6**). Reachability constraints altered target selection in 6 of 14 CU participants, replacing the highest-burden parcel with the highest-burden parcel accessible from the scalp. Illumination orientation was continuously optimized using multi-start Adam ascent with ring-distributed initializations (**Fig. 3c-d**). Optimization trajectories were visualized as both beam-entry paths on participant-specific orthogonal planes and trajectories through the angular dose landscape with two initialization strategies, and the initialization at the nearest International 10–20 scalp positions improved convergence compared with vertex initialization. In the illustrative AD and CU participants, nearest-electrode initialization reached the best solution within the third and first restarts respectively. The deployed angles lay 2.66° and 4.70° angular distance from independently computed MC search LF optima, delivering 88.6% and 91.0% LF of the MC at the optimum, respectively. Individual optimization landscapes of the entire cohort are shown in **Fig. S7**. In the 14 AD participants, optimization increased target LF by a median of 3.8-fold over the best 10–20 scalp positions and recovered 91% of the MC optimum (**Fig. 4a**). In the 14 CU participants, the corresponding gain was 3.9-fold over the nearest 10-20 positions, with 90% recovery of the MC optimum (**Fig. 4b**).

**Fig. 3:**
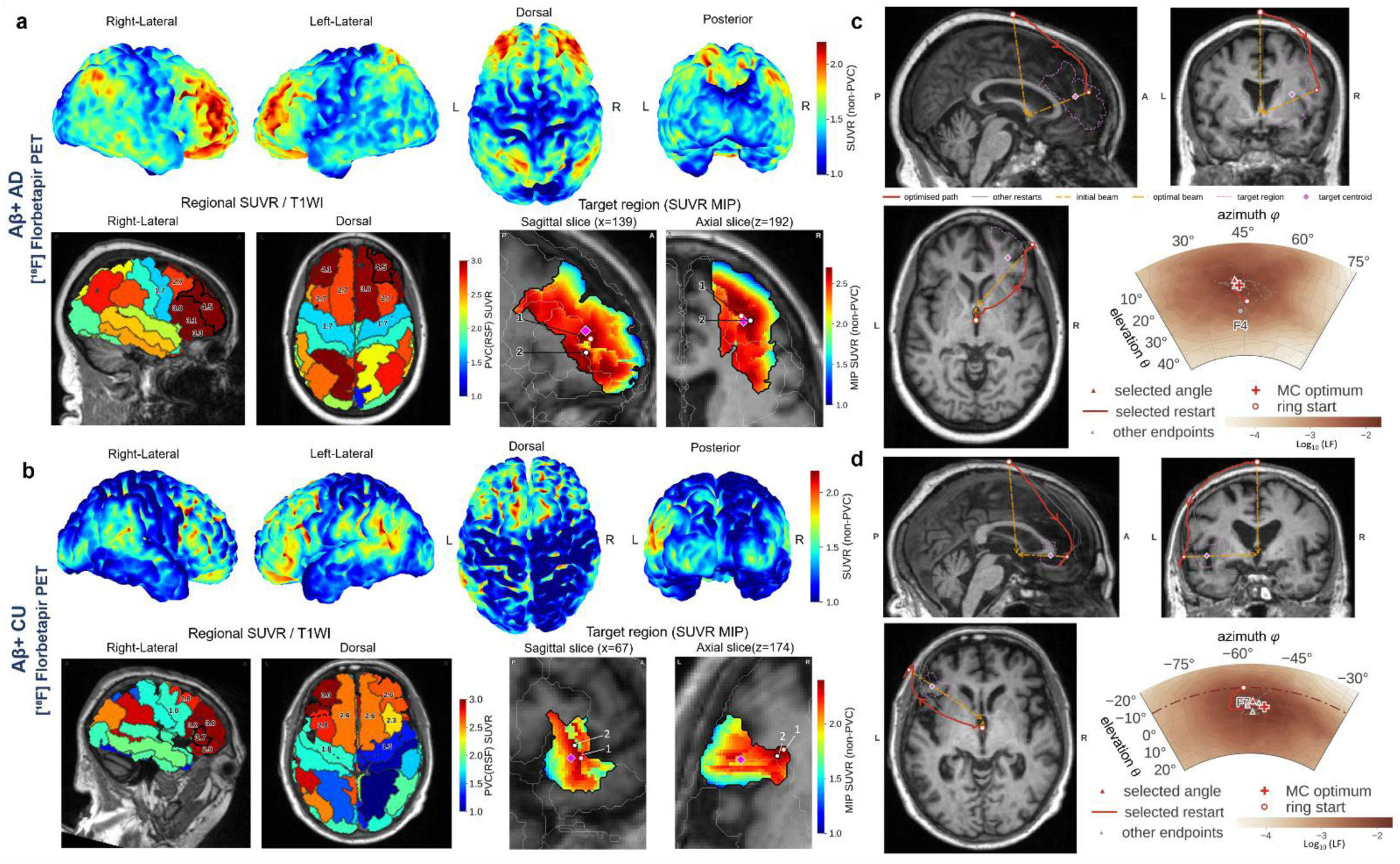
Inverse optimization of illumination guided by Amyloid-PET images. (a-b) Volume of interest (VOI) and target voxel localization definition: (**a**) stimulation target of one Aβ+ AD participant (Centiloid = 111, ID: oas30812). Top, voxelwise SUVR (tracer: [¹⁸F]florbetapir; without partial-volume-correction, i.e. non-PVC) sampled into the cortical ribbon and rendered on the participant’s cerebral surface. Bottom, regional spread function (RSF) PVC-SUVR overlaid on DKT parcellation map and projected over the participant’s T1w image; The target VOI (right rostral middle frontal gyrus) is outlined in black, with the inset showing a maximum intensity projection (MIP) of voxel-wise, non-PVC SUVR within the VOI. Diamond marker, the SUVR-weighted centroid used as the optimization target; circle markers, the voxels with the highest and the second highest SUVR value. (**b**) stimulation target of one Aβ+ CU participant (Centiloid = 85, ID: oas30841). The target VOI, left pars triangularis. (**c-d**) Subject-specific, pathology-informed illumination optimization in the Aβ+ participant. Optimization trajectories of (**c**) the Aβ+ AD participant (ID: oas30812) and (**d**) the Aβ+ CU participant (ID: oas30841) initialized from the Cz site are projected on the sagittal, coronal, and horizontal planes of their own T1w images, with the target VOI and the SUVR-weighted centroid indicated by purple dashed lines and diamond markers; Optimization trajectories initialized from the nearest 10-20 stimulation sites using the same optimizer and objective function are illustrated using a polar map of MNI space. Abbreviations: Aβ, amyloid-beta; AD, Alzheimer’s disease; CU, cognitively unimpaired; DKT, Desikan-Killiany-Tourville; MNI, Montreal Neurological Institute; SUVR, standardized uptake value ratio.

**Fig. 4:**
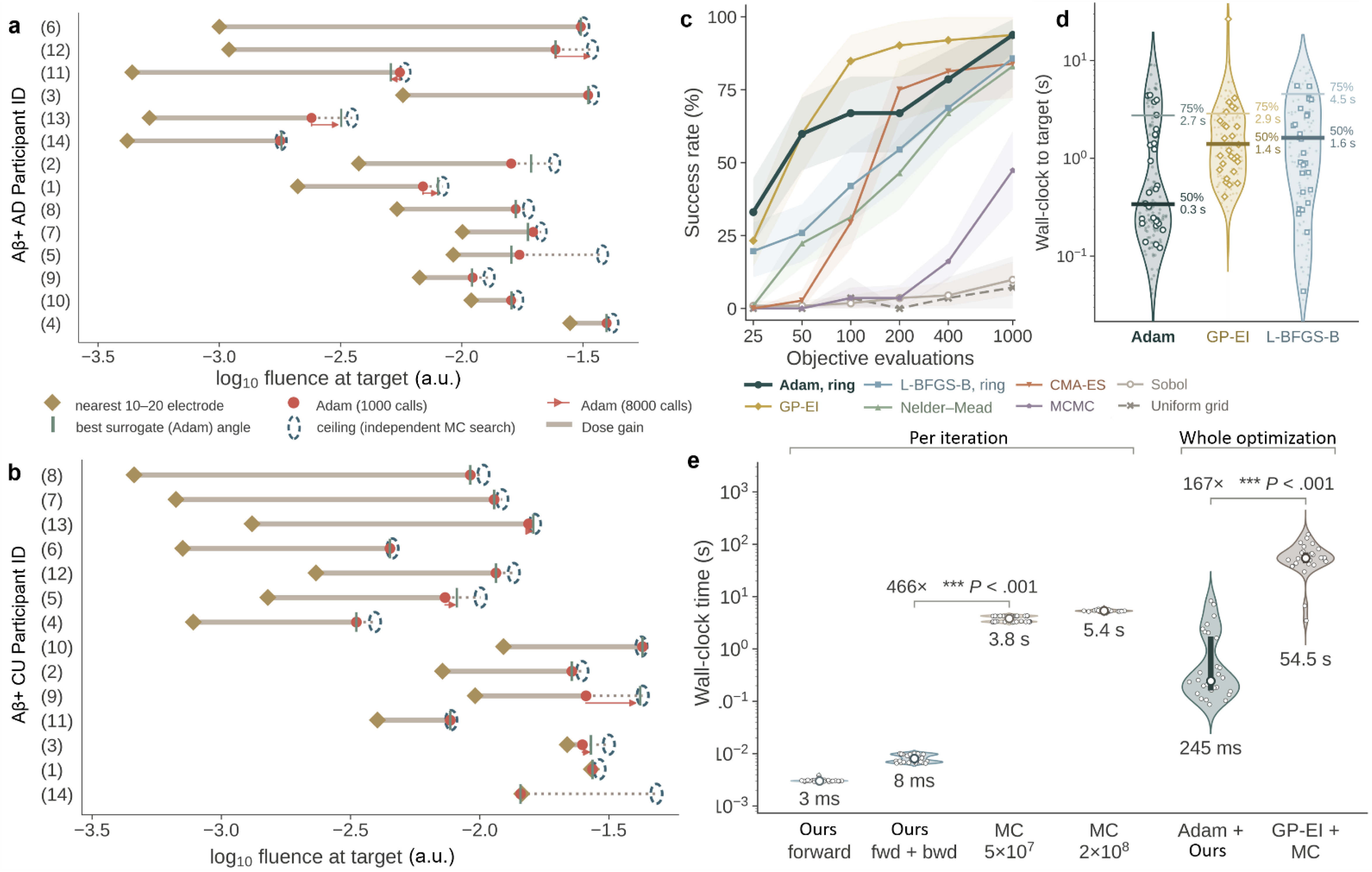
Performance evaluation of PhomiNeuro-driven optimization task. (**a-b**) Light fluence delivered by the optimized illumination parameters at the target VOIs compared to initial setup and the MC optimum: one row per participant, (**a**) Aβ+ AD (n = 14) and (**b**) Aβ+ CU (n = 14). Light fluence gain is calculated from its initial value at participant’s nearest 10–20 stimulation site to the optimized angle predicted by a single 1,000-call ring-restart Adam optimizer, which is posteriorly validated using a 8,000-call Adam optimizer and the MC grid search as ground truth. (**c**) Reliability of eight optimizers on the differentiable objective. Success is a run whose best objective lands within 0.05 decade of the densely searched optimum for that participant; each point pools 28 participants × 4 repeats = 112 runs. Shading, 95% intervals bootstrapped over participants. (**d**) Wall-clock time to reach the success criterion, for the three methods exceeding 90% success at 1,000 calls. Faint dots, individual runs; open markers, per-participant medians (the independent sampling unit); thick bar, T₅₀; thin bar, T₉₀. (**e**) Wall-clock cost of one illumination design. The differentiable surrogate is 466× faster than Monte Carlo per iteration and 167× faster over a whole inverse optimization (brackets, paired median fold change; exact paired sign test, Holm-corrected). Violin, kernel density on log₁₀; bar, interquartile range; white marker, median; open circles, the 28 participants. Abbreviations: Adam, adaptive moment estimation; bwd, backward automatic gradient calculation; CMA-ES, covariance matrix adaptation evolution strategy; fwd, forward surrogate; GP-EI, Gaussian-process expected improvement; L-BFGS-B, limited-memory BFGS with bounds; MCMC, Markov chain Monte Carlo; Nelder-Mead, Nelder-Mead simplex method; Sobol, Sobol sequence / quasi-Monte Carlo sampling.

Eight optimization strategies were compared across the 28 Aβ+ participants with four repeats per method and identical evaluation budget (**Fig. 4c-d**). At 1,000 surrogate evaluations, multi-start Adam and Gaussian-process expected improvement both achieved a 93.8% success rate, defined as reaching an objective within 0.05 LF decades of the best participant-specific value. L-BFGS-B, CMA-ES, and Nelder-Mead achieved success rates of 83.0-85.7%, whereas annealed Metropolis sampling reached 47.3% and non-adaptive Sobol and grid searches remained below 10%. Gaussian-process optimization was more evaluation-efficient at 100 calls, but multi-start Adam reached the success threshold faster in wall-clock time (median time: 0.3 s vs. 1.4 s).

The differentiable surrogate reduced the median cost of a combined forward-and-gradient evaluation from 3.8 s with MCX to 8.1 ms, a median 466-fold per-iteration acceleration across participants (IQR 395-562, **Fig. 4e**). Complete participant-specific optimization required a median of 245 ms, compared with 54.5 s for MC-driven Gaussian-process optimization, yielding a median 167-fold end-to-end speedup (IQR 30-392). Both improvements were significant by exact paired sign tests across participants (Holm-corrected P < 0.001). PhomiNeuro therefore reduced inverse design from approximately one minute to sub-second while retaining independent MC validation of the selected illumination parameters. The same objective was further extended to multiple targets with a similar optimization procedure in both the illustrative participants (**Fig. S8**): the two target voxels defined by the SUVR local maxima on the 1-mm linearly interpolated grid (4.90 mm in the CU participant and 7.48 mm apart in the AD illumination parameters). The joint LF gain was 8.1% and 0.2% on the Aβ+ CU and Aβ+ AD respectively achieved by the two-target equal-weight optimization compared to single-target optimization results. These results suggest that an illumination source position can address multiple targets which are located with moderate distance and guided by effective imaging resolution.

### Cross-participant analysis identifies extracranial thickness as the principal correlate of light delivery efficiency

Cross-participant explainability was evaluated in all 84 held-out participants at the full set of 10-20 illumination sites (**Fig. 5a**). First, correlations among demographic, clinical and anatomical variables reproduced the expected clinical and structure-related associations: scalp thickness declined with PET-centiloid (Spearman’s ρ = -0.38, p < 0.01) and amyloid status (ρ = -0.31, p < 0.05); compensatory skull thickening with BPF declining (ρ = -0.34, p < 0.01) accompanied by aging and AD progress^49–51^; sex difference was most significantly associated with intracranial volume (ρ = 0.65, p < 0.001) and scalp thickness (ρ = 0.43, p < 0.001). Mantel analysis then related each variable to absorbed-energy profiles of grey matter, white matter and CSF predicted by PhomiNeuro, calculated based on dissimilarity in the 10-20 stimulation positions. Only extracranial structures were consistently associated with energy deposition: fat thickness yielded Mantel R = 0.320-0.349, scalp thickness R = 0.239-0.329 and skull thickness R = 0.129-0.250, with P ≤ 0.022 in every tissue. The Aβ burden (Centiloid) and Aβ status, dementia severity, cognitive score, BPF, intracranial volume and age all remained at R ≤ 0.20 without statistical significance.

**Fig. 5:**
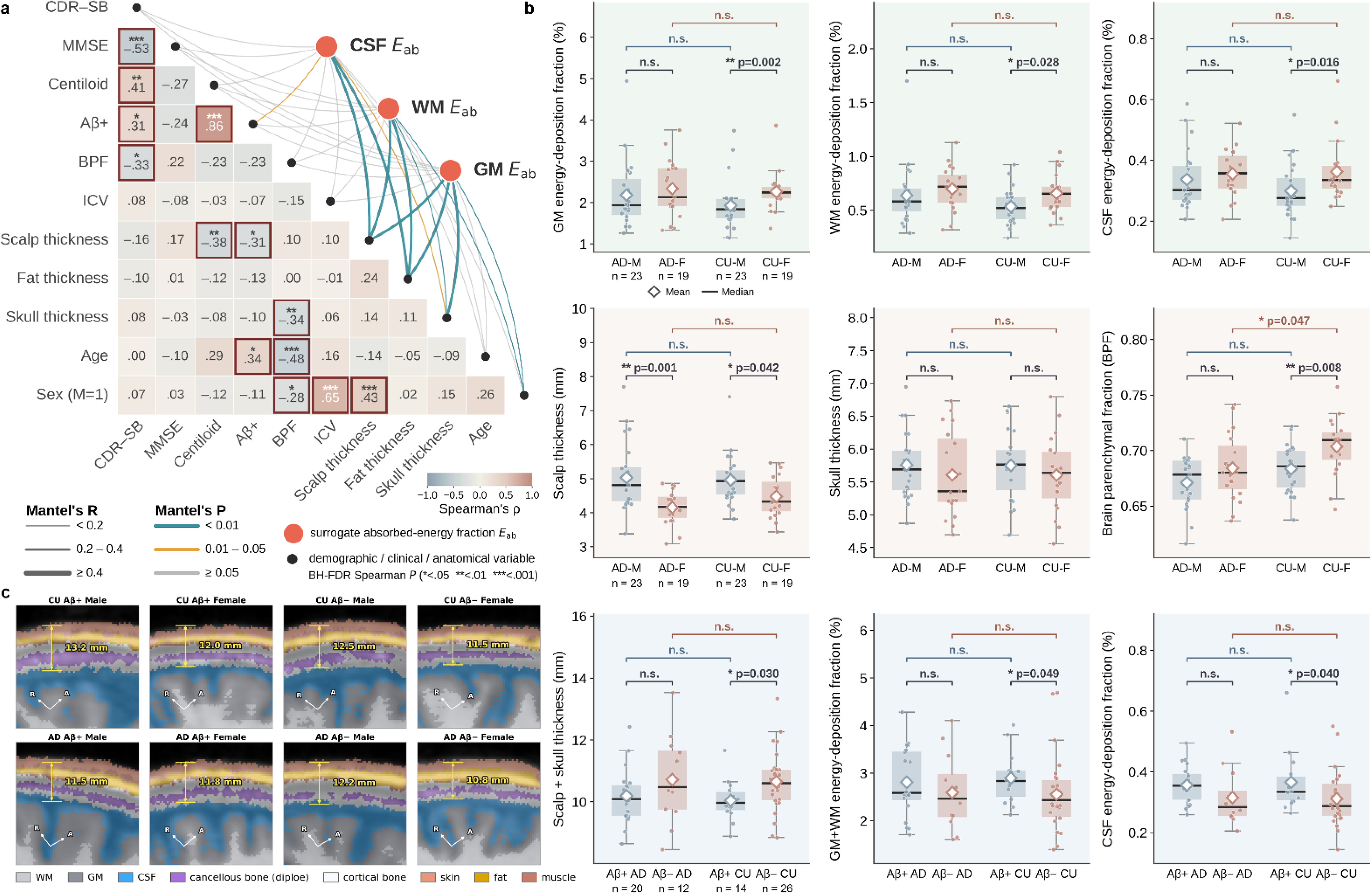
Anatomical and clinical correlates of PhomINR-predicted energy deposition across 84 held-out participants. (**a**) Correlation structure and Mantel correlations across the 84 held-out participants. Cells surviving Benjamini-Hochberg FDR correction at P < 0.05 carry a dark-red border and significance markers. Each chord represents a Mantel test between (i) the dissimilarity matrix of our predicted energy deposition fraction in CSF, white matter (WM) and gray matter (GM) with (ii) the Euclidean distance matrix derived from Pearson correlations of the condensed distance vectors; significance assessed by a two-sided permutation test. (**b**) Comparison of sex and pathology-related differences in energy deposition and anatomical structures, stratified by participants’ sex, diagnosis (n = 84; 42 AD and 42 CN; M:F = 23:19 in each group) and Aβ status of those with accessible Centiloid data (n = 72; Aβ+, Centiloid ≥ 26; Aβ-, Centiloid < 26). Black line and diamond marker, mean and median values over all 10-20 stimulation sites; whiskers, 1.5 × IQR; dots, individual participants. Post-hoc comparisons via two-sided Mann–Whitney U test. (**c**) Scalp and skull thickness at the F4 stimulation site. Eight participants with median scalp-skull thickness were stratified by diagnosis, amyloid status, and sex, whose anatomical structures are displayed via horizontal segmented T1w images. Abbreviations: CDR–SB, Clinical Dementia Rating Sum of Boxes; HC, healthy control; ICV, intracranial volume; MMSE, Mini-Mental State Examination.

Then sex-stratified analyses localized differences in absorbed energy to the CU group (**Fig. 5b, Table S6**). PhomiNeuro-derived absorbed-energy fractions were higher in CU women than in CU men in grey matter (P = 0.002), white matter (P = 0.028), and CSF (P = 0.016), whereas no significant sex differences were observed in AD and no diagnosis differences at fixed sex were detected in any tissue. Independent MC estimates showed the same direction, with significant sex differences in grey matter and CSF. Men had thicker scalps in both diagnostic groups (P = 0.001 and P = 0.042), whereas skull thickness did not differ by sex.

Among the 72 participants with Centiloid data, Aβ+ CU participants had thinner combined scalp-and-skull thickness and greater parenchymal energy deposition than Aβ− CU participants (**Fig. 5b**). At the tissue level, Aβ+ versus Aβ− CU participants showed significantly higher CSF energy deposition (**P = 0.040**) and nonsignificant trends toward higher grey- and white-matter deposition (both P = 0.059; **Table S7**), whereas no corresponding Aβ status-related differences were significant in AD. Among representative participants selected near each subgroup median, combined scalp-and-skull thickness ranged from 10.8 mm in the AD Aβ− female to 13.2 mm in the CU Aβ+ male, revealing a 2.4-mm variation in extracranial optical path length across subgroups (**Fig. 5c**).

We next examined whether the subgroup comparisons results persisted after accounting for correlated covariates. In univariable analyses, absorbed energy decreased with both scalp and skull thickness across all tissues (Pearson r = −0.47 to −0.59; all P < 0.001) but was unrelated to brain parenchymal fraction (|r| ≤ 0.14; all P > 0.2). PhomiNeuro and an independent MC validation agreed in the direction, magnitude, and significance of all nine tissue-by-predictor associations (**Fig. S9**). In multivariable models of the 72 participants with accessible Centiloid data, only scalp and skull thickness remained independently associated with absorbed energy (**Fig. S10**). Standardized coefficients β ranged from −0.40 to −0.56 for scalp thickness and from −0.43 to −0.58 for skull thickness (all P < 0.001), with adjusted R² values of 0.46–0.60. Amyloid burden, BPF, age, sex, and AD-diagnosis were not independently associated with absorbed energy (|β| ≤ 0.19; all P ≥ 0.17). Replacing continuous Centiloid-defined amyloid burden with amyloid status (Aβ+ vs. Aβ-) did not alter these findings: amyloid status remained non-significant for both PhomiNeuro and MC (all |β| ≤ 0.21; all P ≥ 0.23), whereas both thickness measures remained significant (all P < 0.001). Thus, the unadjusted sex- and amyloid-group differences were largely attenuated after accounting for variation in the extracranial tissues, whose structures are the primary drivers of the energy deposition variability at the present clinical cohort.

## Discussion

This study establishes PhomiNeuro as a cross-participant, differentiable light transport surrogate connecting time-resolved individual dosimetry with pathology-guided inverse illumination design. In 84 held-out participants, it achieved leading overall accuracy, spatiotemporal robustness, and cross-cohort generalization, although it did not dominate every benchmark metric. Ablations identified FM-encoded anatomical-optical conditioning as the principal contributor, with domain adaptation, continuous representation, and physics-informed regularization providing complementary contributions. Differentiable querying converted rapid forward prediction into sub-second inverse design, completing participant-specific optimization in 245 ms with an approximately two-order-of-magnitude speedup. Amyloid PET and T1w MR-defined targets and cohort analyses further linked individualized dose planning to pathology and its structural determinants.

Cross-participant generalization arose from three interacting components. The pretrained VISTA3D encoder supplied multiscale anatomical priors, while domain-adaptive fine-tuning aligned them with tissue optical properties that govern photon transport. Source-to-query path integrals summarized illumination paths, and censored hinge supervision stabilized photon-starved deep-tissue regions. The coordinate-conditioned INR then enabled queries at unseen time points and source angles, while the time-dependent diffusion-equation residual constrained the learned field. This residual improved development-test cohort performance and complemented FM encoding, but its removal did not reduce aggregate held-out accuracy; it therefore acted mainly as a physics-informed regularizer rather than the dominant determinant of generalization. The smooth continuous output does not guarantee a more accurate physical solution in deep regions as high-photon MC remained closer to the converged reference in bright LF regions. PhomiNeuro’s advantage is thus continuous, fast, and differentiable for a robust LF prediction, not universal superiority to MC.

Voxel-based MC was selected as the reference because it maps naturally onto cubic head models and tracks photon directions continuously, whereas many deterministic finite-element method (FEM) solutions of radiative transfer require angular discretization; prior comparisons have also reported favorable MC accuracy under matched computational budgets^52^. This fidelity is costly, as spatial accuracy and memory demand depend on voxel resolution, while stable deep-tissue estimates require rapidly increasing photon counts as surviving photon probability decays. Indeed, detecting transmission across an adult head required overcoming approximately 10¹⁸ attenuation and an MC campaign exceeding 850 GPU-hours^53^. A quantum-walk formulation has demonstrated a theoretical quadratic advantage for amplitude estimation, but only for simplified high-energy photon interactions in water, far from participant-specific tissue optics^54^. PhomiNeuro offers a more immediately deployable alternative through amortized inference with a frozen encoder. Although trained using 5 × 10⁷-photon, 2-ns MC simulations tailored to cortical targets, its interpolation in time provided smooth estimates at photon-starved depths, and extrapolated unseen source angles. Training against higher-resolution, higher-photon references may enable resolution-flexible prediction. The full pipeline ran on one 12-GB GPU; chunked queries enabled 8-GB devices, with CPU fallback using approximately 16 GB RAM. Thus, this framework offers high algorithmic scalability and robust cross-platform compatibility on resource-constrained hardware.

Differentiability transformed PhomiNeuro from a forward surrogate into a continuous inverse-design framework, enabling automatic differentiation of target LF with respect to source orientation. Multi-start Adam reached the success threshold in the shortest wall-clock time, whereas Gaussian-process optimization achieved a higher success rate under limited evaluations. The 466-fold per-iteration and 167-fold end-to-end accelerations reduced planning from 54.5 s to sub-second, with gains reaching 31-fold over standard International 10–20 sites. In the representative participant shown in Fig. 3c, the optimal orientation deviated from the direct source-to-target axis, reflecting participant-specific structural and optical heterogeneity across the scalp, skull, CSF, and cortical folds along the transcranial propagation path. One Aβ+ CU test fell below baseline and recovered 29.7% of the MC method, supporting the necessity of multi-start initialization and final MC verification for the optima. As MRI-derived anatomy determined photon transport and PET defined burden-weighted reachable targets, this multi-target design was constrained by 8 mm PET resolution. Close-range target could share one optimized source, whereas separable targets require multiple or sequential sources. Clinical translation will require further validation of dose-response relationships and analysis of multiscale biological effects.

At the cohort level, PhomiNeuro further enabled joint analysis of anatomical, demographic, and pathological factors. In the 84 matched held-out participants, scalp and skull thickness remained the strongest independent correlates. The higher deposition in CU women and Aβ-positive CU participants was attenuated after adjustment, with concordant effects from PhomiNeuro and MC. The observation of subgroup differences only in CU may partly reflect additional heterogeneity in cranial geometry associated with AD, as disease-related regional scalp-to-region distance alterations and age-and sex-dependent variations in skull thickness and scalp-to-cortex distance have been reported^49–51^. Aβ-positive CU individuals may nevertheless retain cognitive resilience through distinct cortical metabolic patterns^55^, providing biological context for amyloid-related differences before overt dementia. No independent amyloid-burden association was detectable at this sample size, warranting larger cohorts. Hemispheric SUVR asymmetry and laterality-dependent tPBM network responses support individualized hemisphere selection potentially using functional MRI (fMRI), fNIRS, and electroencephalography (EEG)^17,56^.

Several limitations define the next steps. Target definition in this study relied on amyloid PET alone. Future extensions should integrate tau-PET, fMRI-defined functional circuits, and tPBM dose-response relationships to enable multidimensional illumination optimization. The framework should also extend beyond single wavelength, prescribed beam profiles, and bounded orientation ranges. Adaptive INR such as MIRE^57^ may better capture multiscale spatiotemporal frequencies. To enhance the fidelity and explainability of interindividual light-transport modeling, future head models could incorporate CT-or zero-echo-time MRI-based skull segmentation and thickness quantification, which identify skull bone more accurately than T1w MRI^58^. Moreover, current MRI-derived models omitted hair and assigned homogeneous literature-based optical properties to each tissue, leaving individual characterization of scalp, skull, hair, and follicular optics unresolved. High-fidelity construction of head models or optical digital twins remains an unresolved upstream module, whereas PhomiNeuro provides a transferable computational core for optical dosimetry and illumination optimization. Clinical translation will further require a clinician-in-the-loop framework for clinician-engineer co-reasoning^59^ that incorporates expert feedback and longitudinal treatment outcomes from AD cohorts.

Looking forward, the PhomiNeuro framework holds strong potential for adaptation to accelerate optical-excitation modeling and predict time-resolved LF fields in pulsed-laser photodynamic therapy^60^, temporal-interference optical stimulation^61^ and LF correction for optical imaging modalities such as transcranial photoacoustic imaging^62–64^. Furthermore, this methodology is poised to extend across species via further domain adaptation, as well as to neuromodulations based on other energy fields such as focused ultrasound stimulation^65,66^. By coupling multimodal MRI/PET imaging with fNIRS/EEG detection and longitudinal treatment-response monitoring, the framework may establish a closed-loop theranostic digital twin that continuously assimilates structural, pathological, functional, and response information to update intervention targets and illumination parameters over time. Therefore, PhomiNeuro provides a highly scalable computational foundation that bridges individualized dosimetry, intervention planning, and response-guided adaptation for precision neuromodulation^67^.

## Methods

### PhomiNeuro Surrogate framework

Our PhomiNeuro surrogate model couples an anatomy-aware FM encoder with a source-conditioned INR network to predict light fuence *Φ* (W·cm^−2^) from subject-specific anatomy, optical properties, illumination parameters, and spatio-temporal coordinates (**Fig. 1a**). The encoder was initialized from a VISTA3D FM variant (NV-Segment-CTMR) pretrained on more than 30, 000 CT and MRI scans covering 345+ anatomical classes^35^. We adapted it to the head model domain using LoRA^36^ with rank 32 and α = 64, together with a masked-autoencoder objective^37^. During adaptation, 64^3^ patches were randomly cropped from the head models with a mask ratio of 0.75 and a patch size of 8^3^. Each head model is represented as a four-channel volume (µ_a_, µ_s_, g, n) at 810 nm and normalized over fixed physical ranges. After adaptation, the encoder was frozen to extract voxel-wise latent space feature vectors^68^ with channel dimensions of [48, 96, 192, 384, 768] (**Fig. S11)**.

For a continuous 4D coordinate (*x*, *y*, *z*, *t*), the local FM-ecoded feature is tri-linearly sampled at (*x*, *y*, *z*) and combined with physics-informed conditioning, including Fourier positional encoding^38^, temporal encoding, source-relative geometric features, i.e., source position, direction, normalized distance, and source-to-point angle. A four-layer GELU MLP with hidden width 512, 512 and 256 then predicts *log*_10_*Φ*(*x*, *y*, *z*, *t*). The source-relative terms provide a geometric prior for distance- and direction-dependent attenuation, while the FM-derived anatomical features encode subject-specfic optical properties and structure. The MLP was trained with a physics-informed multi-loss objective:

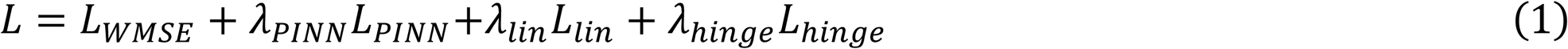

Here *L_WMSE_* is a weighted 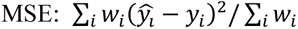 that up-weights near-source tissue layers and CSF layers stratified by fluence decay *d* = log_10_ *Φ_max_* − *y* relative to peak value and by scattering characteristic of the optical media within human head (non-scattering CSF vs. highly scattering tissues). To balance these multi-task objectives, hyperparameter tuning for the λ coefficients was performed exclusively on 20 development-test head models. The resulting final parameters were established as λ_PINN_ = 1.0, λ_lin_ = 0.3, and λ_hinge_ = 1.0. *L_PINN_* is a physics-informed regularization term based on the normalized diffusion residual of the time-dependent diffusion equation (TD DE):

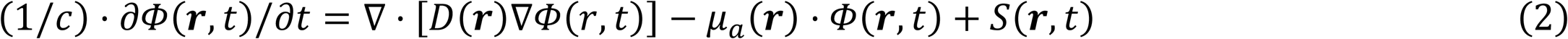

Here *c* is the light speed in tissue (2.188×10^8^ m/s at refractive index *n* = 1.37), ***r*** = (*x*, *y*, *z*), the diffusion coefficient 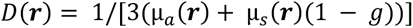. The *L_PINN_* term penalises the residual of the TD DE under three approximations: (1) The residual is only evaluated at non-CSF deep tissue meeting diffusion approximation (decade *d* ≥ 2.5), and thus S(r, *t*) ≈ 0 as the near-source region are excluded; (2) D is treated as locally uniform and ∇ · [*D*∇*Φ*] is approximated as D∇²Φ; (3) The time derivative (1/*c*)*∂Φ*/*∂t* is calculated by (*Φ_j_*_+1_ − *Φ_j_*_−1_)/(2*cΔt*) with Δ*t* = 0.2 ns and time gate j = 1, 2, …, 8. Thus the residual and *L_PINN_* is writen as:

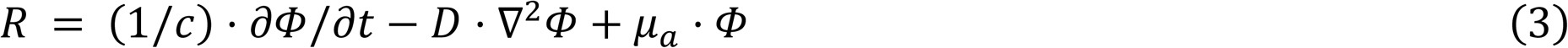

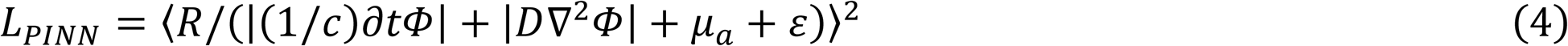

Here ε = 10^−12^.The Laplacian uses a 7-point finite-difference stencil whose half-step (2 mm) is jittered per collocation point. The residual is normalized pointwise by the predicted fluence at the stencil’s center, makes the residual dimensionless and equally weighted across its dynamic range.

*L_in_* is a near-source term to restoring linear fluence fidelity: 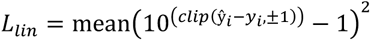 and *L*_ℎ*inge*_ is an one-sided censoring regularization term 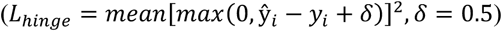 for ***r*** with *d* ≥ 8 at the MC noise floor receiving zero weight in *L*_WMSE_. The loss hyperparameters and model architecture were tuned on the twenty dev-test heads.

By parameterizing continuous fields with separately encoded spatial and temporal coordinates using NeRF-style, octave-spaced sinusoidal Fourier features^38,69^ 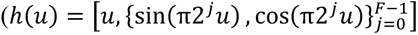, *F* denotes the number of frequency bands; F_s_ = 4 spatial coordinates; F_t_ =3 for the temporal coordinate respecting the Nyquist–Shannon sampling theorem), this INR-based PhomiNeuro Surrogate enables off-grid LF representation and gradient-based inverse design of illumination. Given a target voxel in a volume of interest (VOI) guided by PET image, we optimized the source parameters (*θ*, *φ*) that represent the elevation and azimuth in the MNI spherical coordinate system to maximize the predicted fluence *log*_10_*Φ*. The optimization was based on multi-start Adam gradient ascent, with MCX invoked once for final verification of the optimized source parameters; To evaluate the benefits of differentiability and speed, we further compared our surrogate against other typical gradient-based and gradient-free methods (see method: Inverse optimization of illumination).

### Training and evaluationk

All training and evaluation were conducted on four NVIDIA GH200 GPUs on the high-performance computing (HPC) compute node (i.e. Clariden on Alps) supported by the Swiss National Supercomputing Centre (CSCS) under project ID Prep32 and a0274. The head models from SHARM, ScatterBrains, BrainWeb and OASIS-3 were split at the participant level into training (n = 285), validation (n = 45), development-test (n = 20) and held-out test (n = 84) sets (**Fig. S11a**). All MC simulations derived from a participant remained within one partition; The held-out set was evaluated only once after model selection. The VISTA3D encoder was first adapted through LoRA-based fine-tuning on patches cropped from the training head volumes, with rotations and flips applied for data augmentation. The fine-tuning used AdamW (learning rate: 5×10⁻⁴, cosine decay) with a masked-reconstruction loss; an eight-head spatial redundancy-dropout attention^70^ was leveraged to drop strongly-attended local connections to enforce long-range spatial dependence, as LF integrates the full source-to-point path; a U-Net decoder skip-connected at four scales rebuilt the masked volume. The adapted and frozen encoder was then used to train the INR on 10,800 training LF fields, while 1,810 validation LF fields were used for early stopping and and optimal checkpoint selection. In addition to the standard International 10-20 directions, the training simulations included source directions sampled within their convex hull by Delaunay-based barycentric sampling: 15 directions for each SHARM, BrainWeb and ScatterBrains head and 20 for each OASIS-3 head model.

For the final training of PhomiNeuro’s INR module, supervised coordinates were sampled once and stored in reusable point buffers together with all conditioning features and the ten time gate labels. Each LF fileld contributed K = 3,000 points across the 10,800 training and 1,810 validation LF fields: 45% from the band within six decades of the LF field maximum, 15% from the above-floor bright field and 40% uniformly from tissue. The physics buffer contained 1,296,000 collocation points, each represented by a seven-point finite-difference stencil. The model was trained for 120 epochs with AdamW, an initial learning rate of 10^−3^, weight decay of 10^−5^, cosine learning-rate decay and a batch size of 65,536. The checkpoint with the lowest validation loss was retained.

Evaluation was performed separately on the development-test cohort at all 10-20 source positions (380 LF filelds) and the held-out cohort at Fp1, F7, Cz, T4, Pz and O1 (504 LF filelds). A dense panel comprising all source positions in the held-out participants (1,596 LF filelds) was used only for analyses requiring full angular coverage. Up to 50,000 tissue coordinates were sampled per LF field with deterministic participant-site seeds. A single support mask, defined from the time-integrated MC field at seven decades below the LF maximum, was shared by all ten time gates. Within each gate, the MC reference and prediction were independently truncated at seven decades below their respective maxima, and zero reference values were assigned the corresponding floor. Performance was assessed using the coefficient of determination (R²), Pearson and Spearman correlations, Lin’s concordance correlation coefficient, root-mean-square and median absolute errors, bias, regression slope and variance ratio. Dosimetric agreement was assessed using energy conservation and the gamma pass rate with a 3% amplitude and 2 mm distance criterion over the brightest decade. Off-grid performance was evaluated at spatial and temporal coordinates that did not coincide with the native voxel or time gate centers.

### Benchmark and ablation experiments

The benchmark comprised five volumetric networks (VIT^41^, DynUNet^42^, SegResNet^43^, UNet^44^ and FNO^45^) and two coordinate networks (RFF multilayer perceptron^38^ and SIREN^46^). The volumetric models predicted a fixed ten-channel field, whereas the coordinate models were queried pointwise. All baselines used the same participant partitions, ten-gate targets, output censoring and evaluation protocol as PhomiNeuro and were trained for three independent seeds with default settings from their public implementations.

The ablation matrix contained the complete model and 12 prespecified variants. Input ablations removed the FM feature vectors, replaced the adapted encoder with its unadapted or randomly initialized counterpart, removed the path-integral features, or removed source-relative coordinates. Loss ablations removed the physics residual, linear-space term or censored hinge; weighting ablations replaced decade-stratified weights with uniform weights or set the hinge offset to zero. A joint variant removed both the physics residual and the feature vectors. The radial-monotonicity row was marked not applicable because its weight was zero in the final configuration. Each applicable variant otherwise retained the same data, schedule and hyperparameters. Benchmark models used the K = 3,000 training buffer and were assessed on the held-out cohort. Ablations used a separate K = 1,000 buffer over the same training and validation LF filelds and were assessed on the development-test cohort; results from the two tracks were not pooled.

Inference cost was defined as generation of the complete ten-gate field over all tissue voxels for one illumination LF filelds. Timings were recorded after warm-up with explicit device synchronization and excluded disk input/output. For PhomiNeuro, inference was reported both with the anatomy-dependent feature vectors cached, corresponding to repeated planning for one participant, and with the encoder pass included.

### Inverse optimization of illumination

Inverse optimization was evaluated in 28 amyloid-positive held-out participants with amyloid PET acquired in the same session as the T1w image used to construct the head model (14 participants with AD and 14 CU participants). Cortical parcels were ranked within each hemisphere using regional-spread-function partial-volume-corrected SUVR, and the ten highest-ranked parcels were retained as candidates. Within each candidate parcel, grey-matter voxels were weighted by *max*(*SUVR* − 1, 0) from the uncorrected voxelwise SUVR map registered to the head-model grid. These weights defined the pathology-guided volume of interest, and their weighted centroid defined its representative target coordinate. Optical accessibility was measured along the ray from the anterior commissure through this coordinate to the outer scalp. Candidates with a target-to-scalp depth below 32 mm were considered reachable, and the highest-ranked reachable parcel was selected without substitution.

The illumination pose was parameterized by the elevation θ and azimuth φ of the outward unit vector from the anterior commissure to the source, using the same coordinate convention as the MC simulations. The search domain was continuous; the 19 standard 10-20 positions served as discrete references rather than candidate limits. For target coordinates x_i_ with normalized pathology weights w_i_, the objective was 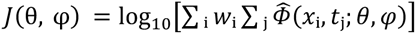, where the inner sum integrates the predicted fluence over the ten simulated gates. Elevation *θ* was constrained to the admissible scalp domain and azimuth *φ* was periodic.

Gradients of *J*(*θ*, *φ*) with respect to *θ* and *φ* were obtained by automatic differentiation through the surrogate. Path-integral features were sampled by differentiable trilinear interpolation; the participant-specific scalp entry radius was tabulated and bilinearly interpolated over the angular domain; and source-dependent features were referenced to the continuous beam-entry point. Analytic gradients were checked against central finite differences using cosine similarity. The adopted optimizer was multi-start Adam ascent with deterministic ring-distributed initializations. It was compared with L-BFGS-B, CMA-ES, Gaussian-process expected improvement, Nelder-Mead, annealed Metropolis sampling, Sobol sampling and a uniform grid at matched budgets of 25, 50, 100, 200, 400 and 1,000 surrogate calls. Four independent repeats were performed for each method, budget and participant. An optimization was considered successful when its best objective was within 0.05 decade of the best value attained for that participant. The deployment configuration used one 1,000-call multi-start Adam run per participant.

Every surrogate-proposed optimum was independently re-evaluated with MC validation and surrogate values were not used to certify the optimized pose. The best standard 10-20 position and the MC-based search reference were computed independently for each participant. Wall-clock comparisons used the same hardware and counted a combined value-and-gradient evaluation as one call for gradient-based methods.

### Study population

All the digital head models were constructed using anatomical model or imaging data of 434 participants under approved data use agreements, including 225 pre-segmented head models: 196 from SHARM models (15 tissue classes, 1 mm isotropic), originating from the IXI healthy adult cohort^71^, 16 normal brain digital phantoms from BrainWeb (11 tissue classes, 0.5 mm isotropic)^72,73^; and 13 from scatterBrains derived from healthy adult MRI (5 tissue classes, 1 mm isotropic)^74^. The remaining 209 participants includes a longitudinal cohort of normal aging and AD, which were obtained from OASIS-3 restricted to sessions with co-acquired T1w data and time-of-flight MR angiogram (TOF-MRA) for brain tissue and cerebrovascular segmentation^75^. OASIS-3 also provided a clinically phenotyped cohort (n = 84) for this study (**Table 2**), serving as the held-out dataset of participants clinically diagnosed with AD (n = 42, CDR ≥ 0.5) and 42 cognitively unimpaired (CU) controls (n = 42, CDR = 0) matched pairwise on age and sex (77.0 ± 7.6 vs. 77.1 ± 7.7 years; famle: male = 19:23). Neither diagnostic or demographic labels were used during the training stage. Clinical and molecular annotation was obtained, including NACC Uniform Data Set Form D1 diagnoses, the CDR value and its Sum of Boxes, the Mini-Mental State Examination (MMSE), APOE genotype and demographics; Amyloid burden was quantified with the OASIS-3 PET Unified Pipeline (PUP)^75^. The cohort contains two tracers, [¹⁸F]florbetapir and [¹¹C]Pittsburgh compound B (PiB). Therefore, the global burden was expressed on the tracer-harmonized Centiloid scale and Aβ positivity (Aβ+) was defined as Centiloid ≥ 26, a threshold supported by the latest clinical research conclusions^47,48^. Of the 72 participants with amyloid PET data, 20 of 32 AD patients were Aβ+ and 14 of 40 CU participants were Aβ+.

**Table 2:**
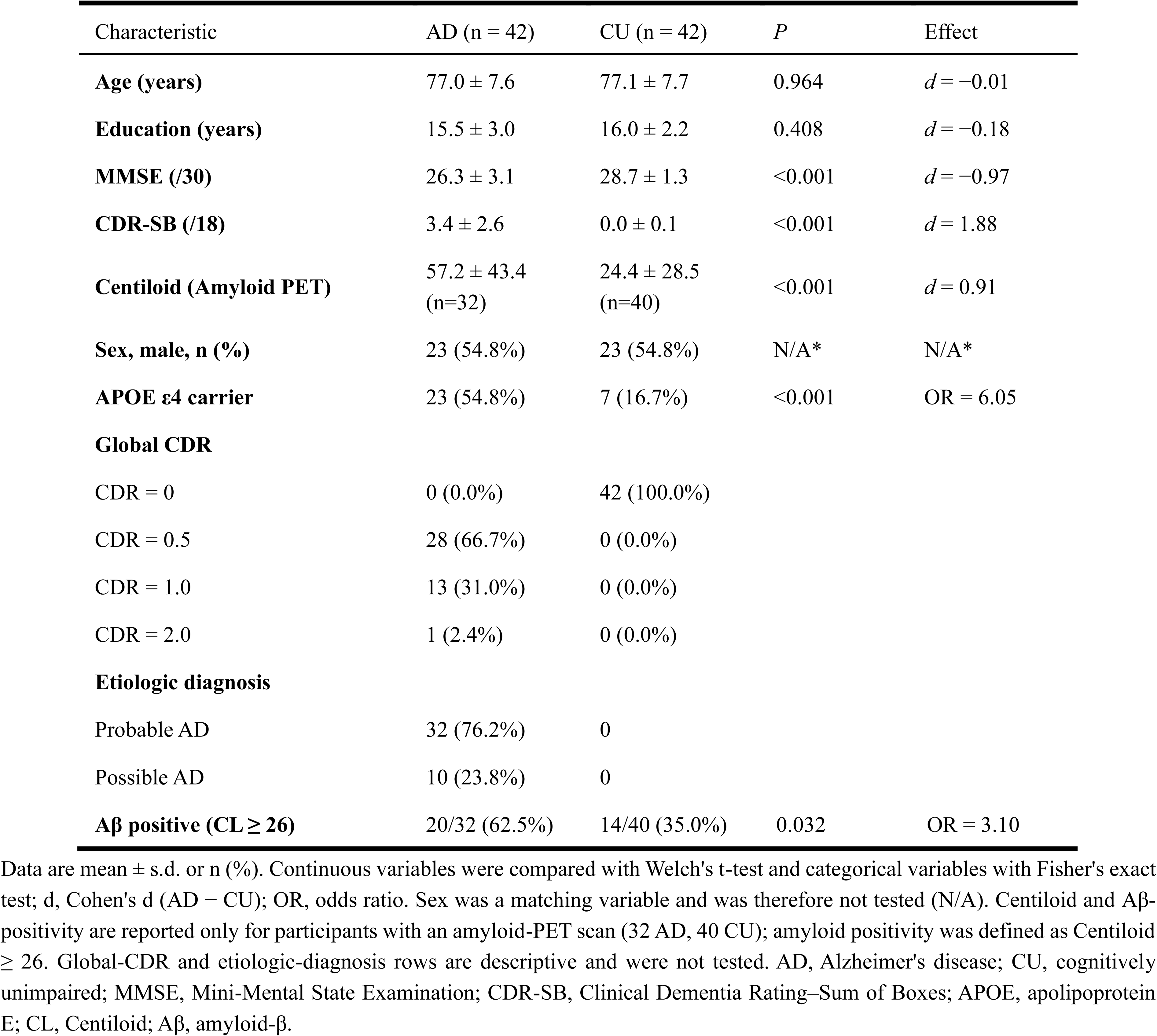
Characteristics of the held-out OASIS-3 test cohort.

| Characteristic | AD (n = 42) | CU (n = 42) | *P* | Effect |
|---|---|---|---|---|
| **Age (years)** | 77.0 ± 7.6 | 77.1 ± 7.7 | 0.964 | *d* = −0.01 |
| **Education (years)** | 15.5 ± 3.0 | 16.0 ± 2.2 | 0.408 | *d* = −0.18 |
| **MMSE (/30)** | 26.3 ± 3.1 | 28.7 ± 1.3 | <0.001 | *d* = −0.97 |
| **CDR-SB (/18)** | 3.4 ± 2.6 | 0.0 ± 0.1 | <0.001 | *d* = 1.88 |
| **Centiloid (Amyloid PET)** | 57.2 ± 43.4<br>(n=32) | 24.4 ± 28.5<br>(n=40) | <0.001 | *d* = 0.91 |
| **Sex, male, n (%)** | 23 (54.8%) | 23 (54.8%) | N/A* | N/A* |
| **APOE ε4 carrier** | 23 (54.8%) | 7 (16.7%) | <0.001 | OR = 6.05 |
| **Global CDR** |  |  |  |  |
| CDR = 0 | 0 (0.0%) | 42 (100.0%) |  |  |
| CDR = 0.5 | 28 (66.7%) | 0 (0.0%) |  |  |
| CDR = 1.0 | 13 (31.0%) | 0 (0.0%) |  |  |
| CDR = 2.0 | 1 (2.4%) | 0 (0.0%) |  |  |
| **Etiologic diagnosis** |  |  |  |  |
| Probable AD | 32 (76.2%) | 0 |  |  |
| Possible AD | 10 (23.8%) | 0 |  |  |
| **Aβ positive (CL ≥ 26)** | 20/32 (62.5%) | 14/40 (35.0%) | 0.032 | OR = 3.10 |

Data are mean ± s.d. or n (%). Continuous variables were compared with Welch's t-test and categorical variables with Fisher's exact test; d, Cohen's d (AD − CU); OR, odds ratio. Sex was a matching variable and was therefore not tested (N/A). Centiloid and Aβ-positivity are reported only for participants with an amyloid-PET scan (32 AD, 40 CU); amyloid positivity was defined as Centiloid ≥ 26. Global-CDR and etiologic-diagnosis rows are descriptive and were not tested. AD, Alzheimer's disease; CU, cognitively unimpaired; MMSE, Mini-Mental State Examination; CDR-SB, Clinical Dementia Rating–Sum of Boxes; APOE, apolipoprotein E; CL, Centiloid; Aβ, amyloid-β.

Defining amyloid targets at the voxel level requires exact spatial correspondence between the molecular map and the digital head model. This analysis was therefore restricted to participants for whom PUP-processed amyloid PET was available from the same imaging session as the T1w scan used to build their head model. Of the 34 Aβ+ participants in the held-out cohort, 28 met this criterion; the six exclusions were confined to the AD group and were attributable to data availability (two had no PUP output, and four had PUP not concurrent with the T1). The resulting subgroups are 14 Aβ+ AD participants (Centiloid 40.1 to 130.7; 13 imaged with [¹⁸F]florbetapir, one with[¹¹C]PiB) and 14 Aβ+ CU participants (Centiloid 26.3 to 92.5; all [¹⁸F]florbetapir). These 28 participants supply the voxel- and parcel-level amyloid targets for the illumination optimization task evaluated later.

### Data preprocessing pipeline

All 434 head models were rigidly registered to an 1 mm MNI152 template^76^ and resampled onto a common grid of 224 × 256 × 300 voxels, with the anterior commissure (AC) located at voxel (112, 148, 175), corresponding to the spatial origin [0, 0, 0] mm of the MNI coordinate system. Registration was multi-started from two centerd initializations to retain transform with the highest Mattes mutual information using SimpleITK^77^. Tissue labels were resampled by nearest neighbor and then mapped to optical properties (absorption coefficient, scattering coefficient, anisotropy factor and refractive index) compiled to model light transport in human head models during the tPBM treatment using near-infrared light with a wavelength of 810 nm^7,15^, using the optical property values reported in the reference^78^. OASIS-3 tissue labels were derived here from the T1w scan and the TOF-MRA of the same imaging session, all acquired at 3T under the OASIS-3 imaging protocol^75^; The T1 was resampled to a 1 mm isotropic canonical grid, and a head mask was formed by intensity thresholding, morphological closing and hole filling. This mask defines the scalp-air boundary at which illumination enters the head model, and bounds the bias-field correction, the extracranial labelling and the vessel segmentation. Intensity inhomogeneity within the mask was corrected by N4 bias field correction^79^; The brain was extracted with deepbet, a convolutional brain-extraction network^80^, and CSF grey matter and white matter were segmented within the resulting mask using the ANTs Atropos expectation-maximisation classifier with a Markov random field prior^81^. Skull, skin, fat, muscle, and sinus air were segmented with GRACE, a whole-head segmentation network trained on older adults^82^; Head-mask agreement was verified for each participant (Dice 0.97–0.98). Brain classes were retained from Atropos, whose CSF estimate is comparatively liberal in this elderly cohort; as the absorption coefficient of CSF is roughly one-tenth that of grey matter, this choice has an outsized effect on simulated photon transport. Intracranial vessels were segmented from the angiogram, mapped onto the T1 grid analytically through the shared scanner coordinate frame; voxels above the 99.3rd intensity percentile were labelled as vessel, and connected components smaller than 20 voxels were discarded. The segmentations define nine tissue classes: air, skin, skull, CSF, grey matter, white matter, vessel, fat and muscle and the scalp, skull, and fat thickness was measured using the BrainCalculator pipeline^50^.

For the 28 Aβ-positive held-out head models involved in the optimization task, amyloid PET data paired with T1w were obtained from OASIS-3 for each participant, including (1) a voxelwise SUVR image without partial-volume correction (PVC) and (2) a lookup table of regional SUVR with regional spread function (RSF) PVC, together with a FreeSurfer parcellation map^83^. The voxelwise SUVR image and the parcellation map were supplied on a 256³ FreeSurfer conformed 1.0 mm isotropic grid, which is defined in the same scanner (world) coordinate system as the native T1w image. Then the previously calculated rigid transform for the head model was applied to resample the voxelwise SUVR image and the parcellation map into the MNI coordinate system^84^; Trilinear interpolation was used for the SUVR image and nearest-neighbor assignment for the parcellation map.

For the MC reference simulation dataset of LF delivered by each NIR brain stimulation LF filelds, the illumination positions were standardized to the 19 sites of the International 10–20 system for EEG electrode positioning in the MNI space^85^ (**Fig. S10b**), with unit direction vectors for the 19 sites derived from the standard 10–20 electrode coordinate. All heads rigidly registered to MNI with the anterior commissure (AC) fixed as the common origin of the coordinate; each beam has a direction from its stimulation site to AC, with the illumination source 15.0 mm outside the scalp surface. Source position was encoded by elevation (*θ* = arcsin(*n_z_*) ∈ [−*π*/2, *π*/2]) and the azimuth (*φ* ∈ (−*π*, *π*]) is uniquely defined by (sin *φ* = *n_x_*/ cos *θ*) and (cos *φ* = *n_y_*/ cos *θ*) . of the outward unit vector *n̂* = [*n_x_*, *n_y_*, *n_z_*] from AC to the stimulation site. This parameterization ensured anatomical correspondence across subjects. Reference LF fields were computed via MCX^23^, a GPU MC photon transport solver (810 nm, 5×10⁷ photons per LF fileld, 1 mm grid, 5-mm radius disk source, impulse excitation with normalized total energy, 2-ns window divided into ten 0.2-ns gates).

### Statistical analysis

Continuous clinical variables are reported as mean ± s.d. and were compared between the AD and CU groups using Welch’s t-test; categorical variables were compared using Fisher’s exact test. Effect sizes were reported as Cohen’s d for continuous variables and odds ratios for categorical variables. Sex was not tested in the baseline comparison because it was a matching variable. The non-normally distributed MMSE, CDR Sum of Boxes and Centiloid measures were additionally compared using the two-sided Mann-Whitney U test. Amyloid positivity was defined at each analysis as Centiloid ≥ 26; participants without an amyloid scan were treated as missing rather than amyloid-negative.

Model-level metrics were reduced in a fixed field-to-gate-to-seed hierarchy. Within each gate, field-level values were averaged except for ratio-type quantities, including absorbed-energy ratio, regression slope and variance ratio, which were summarized by the median. The ten gate values were then averaged. Three training seeds yielded the reported mean and sample s.d.; distributions across participants are reported separately as median and interquartile range. A difference between two three-seed models was considered larger than reinitialization variability when it exceeded 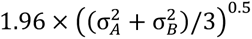^5^, where σ and *σ* denote their across-seed s.d. For comparison with a single-seed arm, the corresponding band was 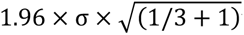 where *σ* denotes the s.d. of the three-seed model. These bands were used as descriptive stability criteria and were not interpreted as hypothesis tests. Associations between participant characteristics and spatial absorbed-energy profiles were evaluated using two-sided Mantel tests.

For each tissue, Bray–Curtis dissimilarities were calculated between participants’ predicted absorbed-energy-fraction profiles across the International 10–20 stimulation positions. These were correlated with pairwise Euclidean distances in each clinical or anatomical variable using Pearson’s Mantel *r*. Interparticipant variability in grey-matter energy deposition was examined by multivariable linear regression including amyloid status, scalp thickness, skull thickness, brain parenchyma fraction, age, sex and diagnosis. Continuous variables were standardized before fitting, and effects are reported as standardized β coefficients with 95% confidence intervals. Identical models were fitted to PhomiNeuro and MCX estimates, and group-level interpretations required concordant effect directions. Omnibus comparisons across more than two groups used the Kruskal-Wallis test for continuous variables and the Fisher-Freeman-Halton exact test with MC permutation for categorical variables. Crossed designs with unequal group sizes were analyzed by two-way analysis of variance with type II sums of squares, with interaction terms reported.

All hypothesis tests were two-sided. Benjamini-Hochberg false-discovery-rate correction was applied to the correlation matrix in Fig. 5a and to the 34 parcel-wise asymmetry tests within each cohort cell in Fig. S5b (q < 0.05). Holm correction was applied to the exact paired sign tests for the wall-clock comparisons in Fig. 4e. All remaining analyses were exploratory and uncorrected. Statistical analyses were performed in SciPy and statsmodels; permutation procedures used fixed random seeds.

## Data availability

The anatomical data and head models analyzed in this study were obtained from openly available or controlled-access research resources. The SHARM segmented head models are available from Figshare (https://doi.org/10.6084/m9.figshare.31150387) and were derived from the IXI MR dataset (https://brain-development.org/ixi-dataset/). The BrainWeb anatomical models are available from the McConnell Brain Imaging Centre (https://brainweb.bic.mni.mcgill.ca), and the ScatterBrains head models are available at https://github.com/wumelissa/scatterBrains. OASIS-3 is a controlled-access resource available upon application through OASIS (https://www.oasis-brains.org/access) and was used under the OASIS Data Use Agreement. OASIS-3 data are not redistributed with this paper. Source data underlying the reported results are provided with this paper.

## Code availability

The codes used in this work are publicly available at: https://github.com/dddshzy/PhomiNeuro

## Acknowledgements

This work was supported by a grant from the Swiss National Supercomputing Centre (CSCS) under project ID Prep32 on the Alps (RN). XW, GL, SD acknowledges the National Key Research and Development Program of China (Grant No. 2025YFE0218400), and SD acknowledges the Program of China Scholarship Council (Grant No. 202506010168). Data were provided in part by SHARM derived from the IXI MR dataset and OASIS-3: Longitudinal Multimodal Neuroimaging: Principal Investigators: T. Benzinger, D. Marcus, J. Morris; NIH P30 AG066444, P50 AG00561, P30 NS09857781, P01 AG026276, P01 AG003991, R01 AG043434, UL1 TR000448, R01 EB009352. AV-45 doses were provided by Avid Radiopharmaceuticals, a wholly owned subsidiary of Eli Lilly.

## Author contributions

Conceptualization: SD, GL, YY, RN, XW.

Methodology: SD, MG, LY, WR

Investigation: SD, MG, YY, RN.

Data curation & formal analysis: SD.

Resources: RN, AR, XW.

Writing – original draft: SD.

Writing – review & editing: RN, WR.

Supervision: AR, RN, WR, XW.

