## Supplementary Materials for "Toward pathology-guided illumination optimization for neuromodulation with a foundation-model-encoded light transport surrogate"

Sihan Dong[1,2], Maoye Guan[2], Liqin Yang[2,3], Guoquan Liu[1], Axel Rominger[2], Yixuan Yuan[4], Wuwei Ren[5#], Ruiqing Ni[2#], Xunbin Wei[1#]

[1] *Institute of Medical Technology, Peking University Health Science Center, Biomedical Engineering Center, Institute of Advanced Clinical Medicine, Peking University, Beijing 100191, China.*

[2] *Department of Nuclear Medicine, Inselspital, Bern University Hospital, University of Bern, Bern 3010, Switzerland.*

[3] *Department of Radiology, Huashan Hospital, Fudan University, Shanghai 200040, China*

[4] *Department of Electronic Engineering, The Chinese University of Hong Kong, Hong Kong, China*

[5] *School of Information Science and Technology, State Key Laboratory of Advanced Medical Materials and Devices, ShanghaiTech University, Shanghai 201210, China*

**The Supplementary Materials include:**

- Fig S1 to S11,
- Table S1 to S7,
- Supplementary Video S1 and S2.

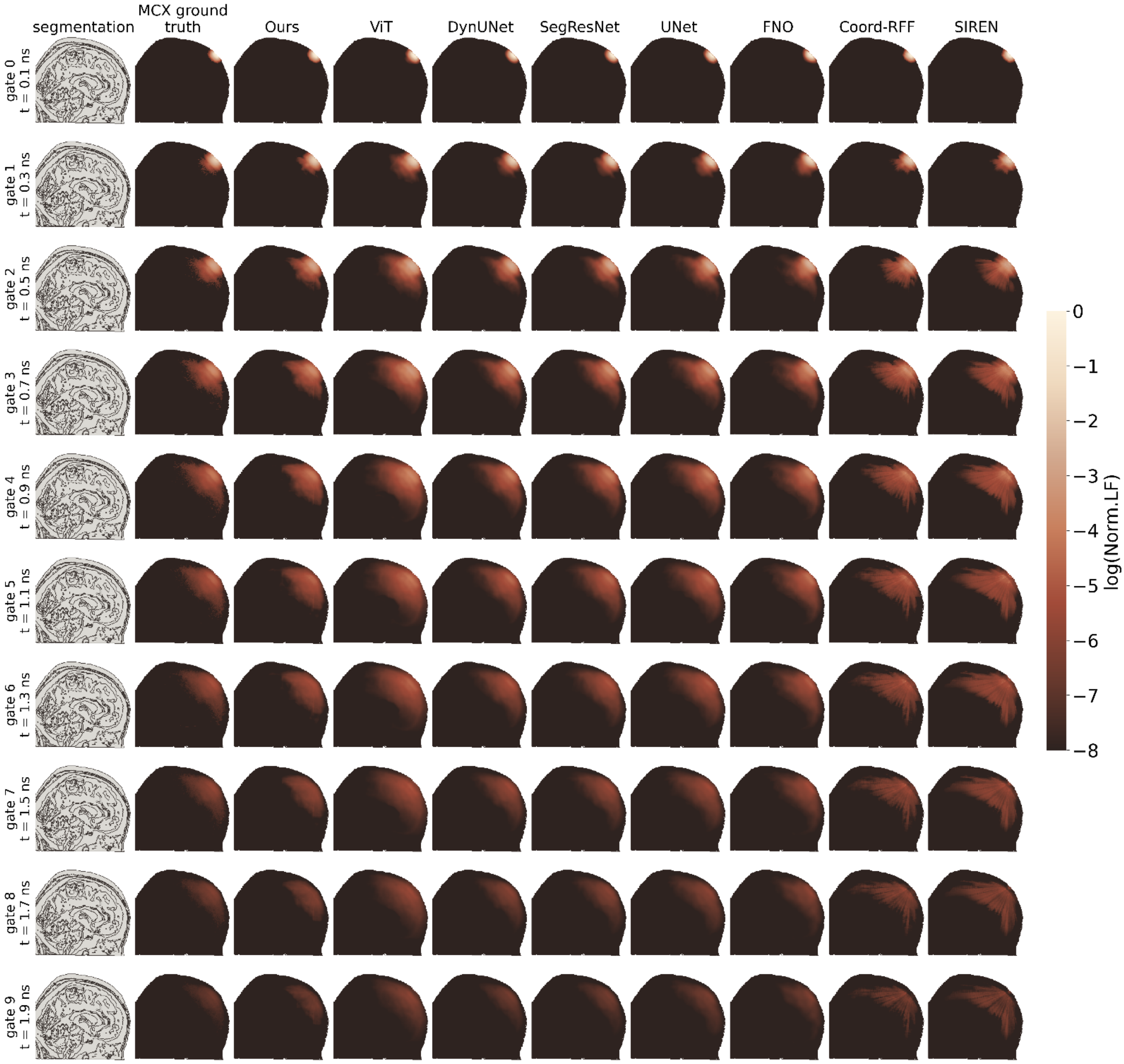


**Fig. S1: Time-resolved light fluence fields on a development-test head.** Light fluence in the plane through the beam (illumination via the Fz stimlation site), one row per 0.2 ns gate. Columns: tissue segmentation, Monte Carlo ground truth calculated by MCX, ours, and seven baseline surrogates.

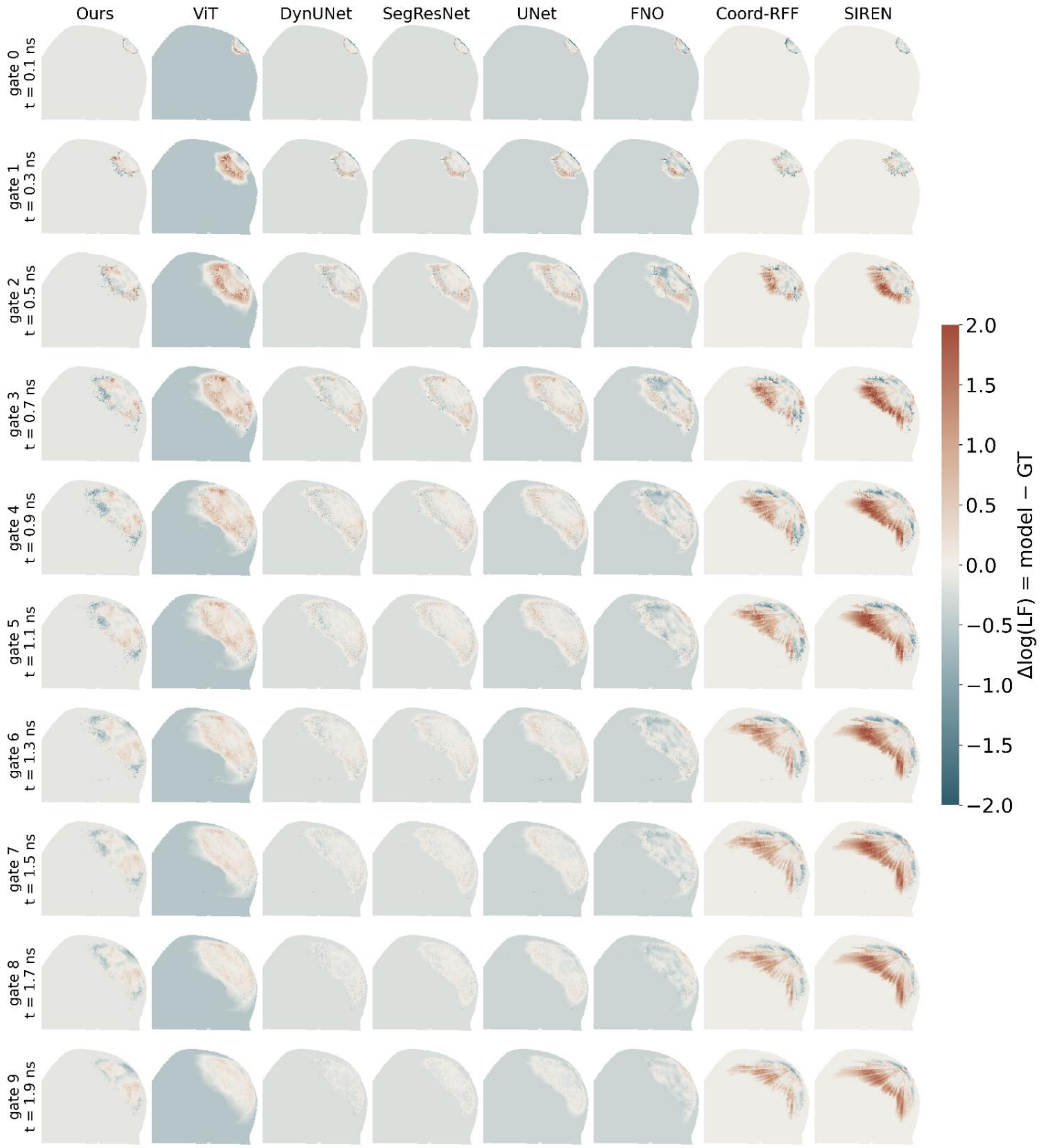


**Fig. S2: Prediction residual of each surrogate network against the Monte Carlo ground truth.** Signed residual, Δlog(LF) = model prediction− ground truth, is computed per head model for the same gates and networks as in Fig S1. Each LF field is floor-clamped ($\log(LF)_{max}$-8) prior to differencing, and residuals are evaluated over the entire volume. LF, light fluence.

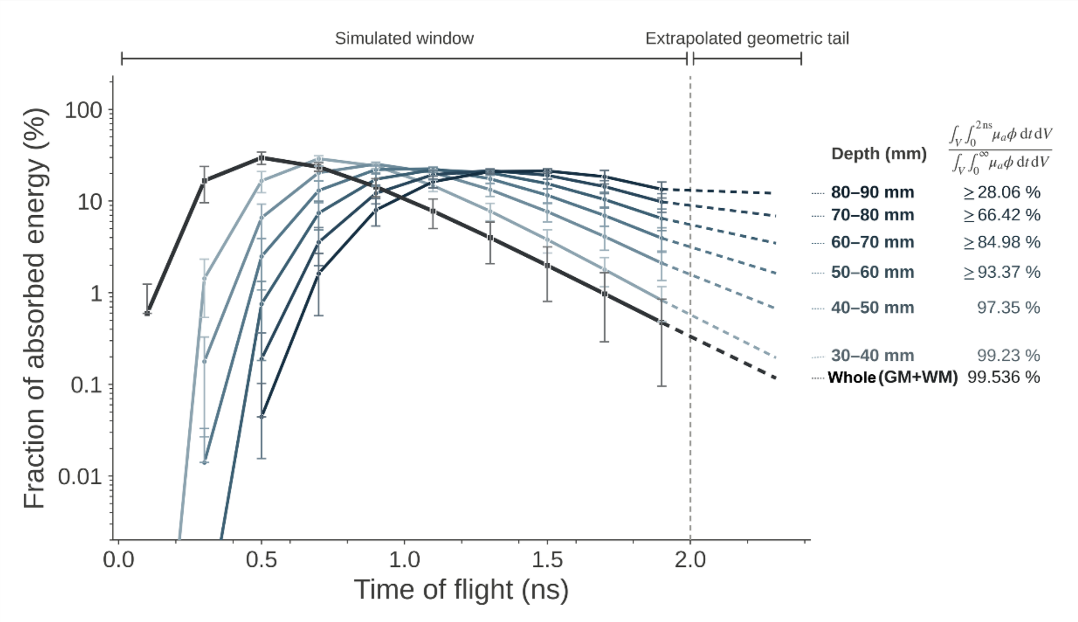


**Fig. S3: Fraction of absorbed energy captured by the simulated 2 ns window in brain parenchyma.** Time-resolved absorbed-energy fractions across penetration-depth bands and the whole brain parenchyma (GM and WM). Curves display cumulative percentages, defined by the 0–2 ns to 0–∞ absorbed-energy integral ratio. Solid lines, the simulated window; dashed lines, extrapolated geometric tails beyond 2 ns; dashed vertical line, the 2 ns window edge; Error bars, s.d. across the cohort.

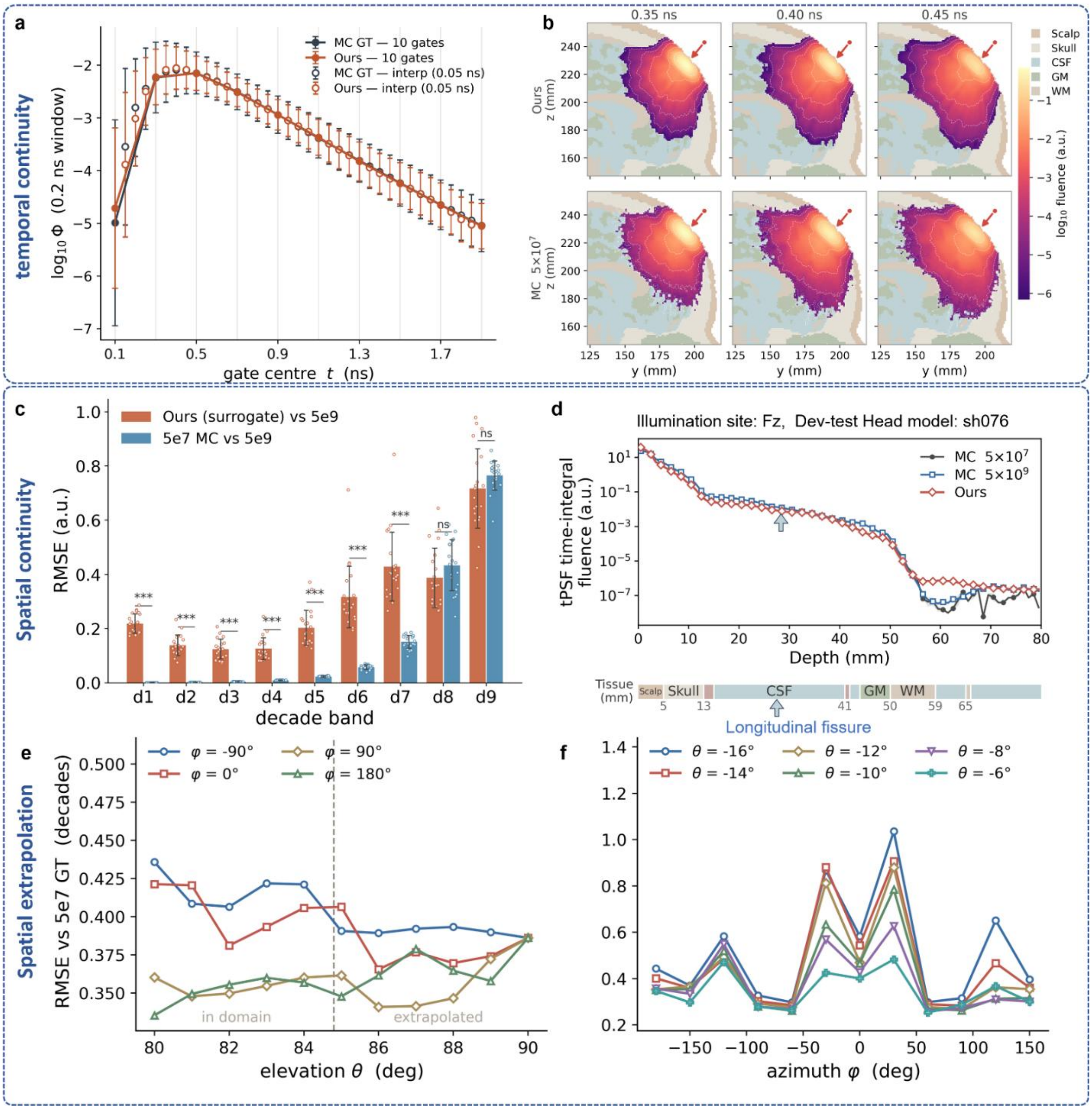

**Fig. S4: Off-grid temporal and angular generalization.** (**a**) Interpolation of temporal point spread function (tPSF) of light fluence at first gray matter (GM) voxel on beam axis under Fz (average over twenty dev-test head models). Ground truth (GT): 40-gate $5\times10^7$ MC. (**b**) Cross-sectional interpolation. Sagittal light fluence field (head model ID: sh076) at off-grid instants (0.35, 0.40, 0.45 ns). Top: PhomiNeuro; bottom: $5\times10^7$ Monte Carlo (MC). PhomiNeuro delivers smooth fields where MC full into speckled noise. (**c**) Accuracy by decade band. RMSE against $5\times10^9$ reference MC GT across nine bands. Points: 20 dev-test head models; error bars: s.d.; significance: paired Wilcoxon signed-rank test.(**d**) Integrated light fluence versus depth. Single head model under Fz stimulation site. The $5\times10^7$ baseline suffers photon starvation beyond 50 mm; PhomiNeuro and $5\times10^9$ MC remain stable.(**e**) Over-elevation extrapolation. Fluence error against $5\times10^7$ MC for θ = 80–90° (training limit: 84.8°). Curves: four azimuths φ.(**f**) Under-elevation extrapolation. As in (**e**) for θ = −6° to −16° (out-of-range).

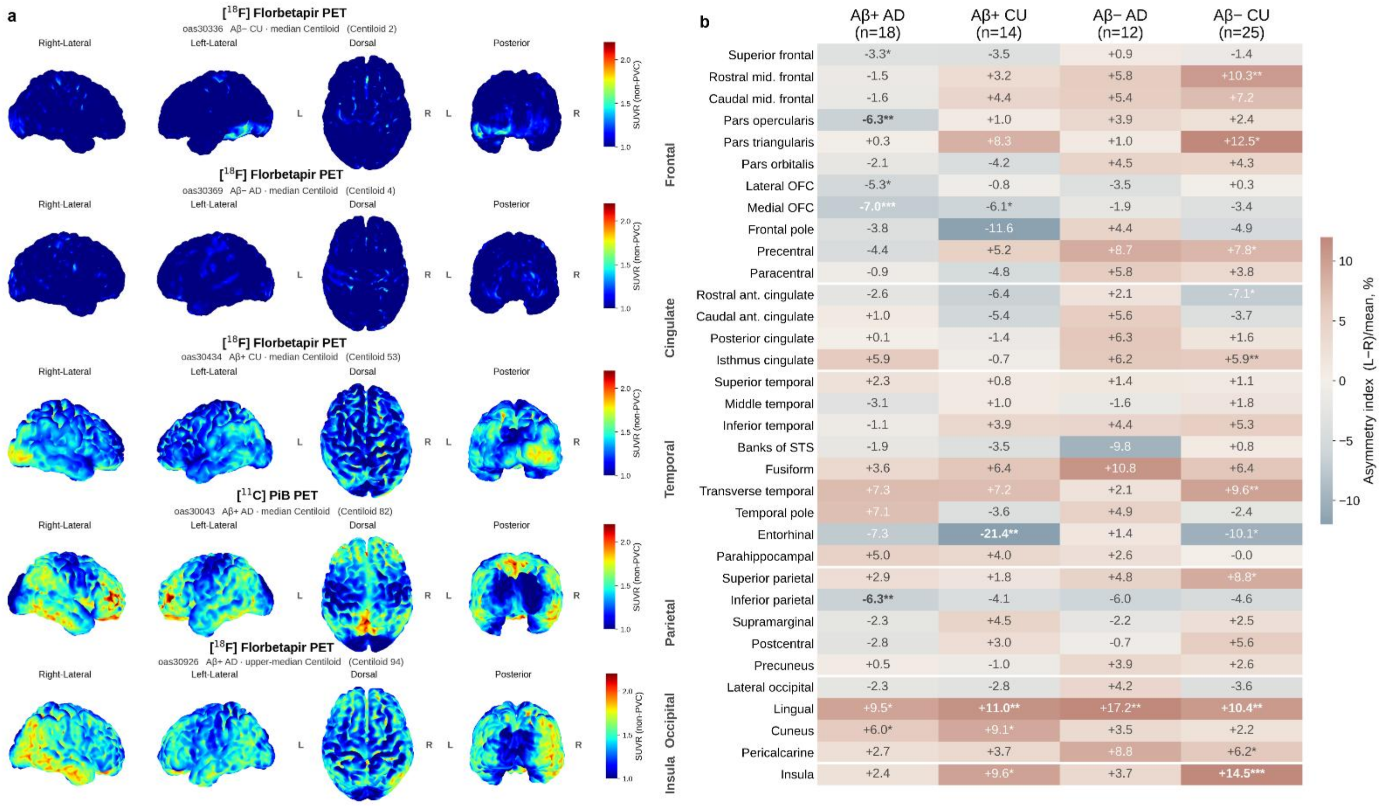


**Fig. S5: Amyloid distribution and hemispheric asymmetry in the held-out OASIS-3 cohort.** (**a**) Cortical SUVR renderings of five representative participants (median Centiloid per cohort). Participant IDs: oas30336, oas30369, oas30434, oas30926 ([$^{18}$F]florbetapir); oas30043 ([$^{11}$C]PiB). Cohorts are stratified by diagnosis (AD/CU) and amyloid status (Aβ+/Aβ−). (**b**) Left-right asymmetry of cortical SUVR across cohorts (n = 18, 14, 12, 25). Heatmap and values indicate mean asymmetry (red: left > right). Stars: nominal significance (within-group paired Wilcoxon signed-rank test); bold: FDR-corrected significance (Benjamini–Hochberg q < 0.05). Abbreviations: SUVR, voxelwise standardized uptake value ratio; Aβ, amyloid-beta; AD, Alzheimer's disease; CU, cognitively unimpaired;

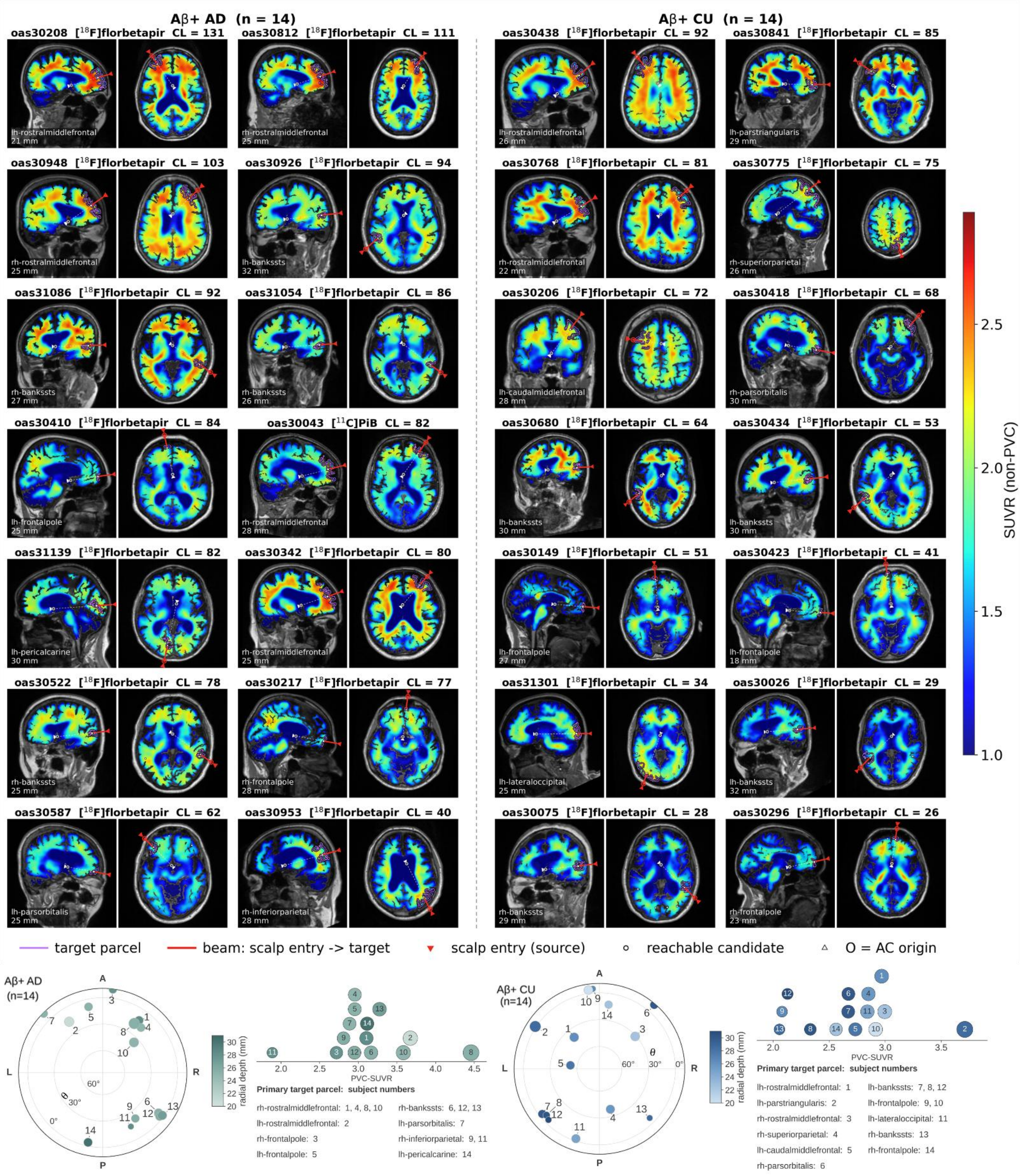


**Fig. S6: Primary stimulation targets in the amyloid-positive (Aβ+) AD and CU cohorts. Top**: target parcel structure and beam orientations overlay. Beam directions (red) and primary target contours (purple) are overlaid on non-PVC PET and T1w MRI for 28 participants (14 AD, 14 CU), ordered by Centiloid. Red/white circles: horizontal projections of scalp beam entry and anterior commissure. **Bottom**: cohort-specific target positions, depths, and amyloid burden. Polar maps display 14 targets per cohort; marker size encodes partial-volume-corrected SUVR (axis beneath); color encodes radial depth. Target parcel identities are listed below. For the Aβ + AD cohort, the OASIS-3 individual IDs (with the "oas3" prefix omitted) and their corresponding single-target optimization task IDs are 0522 (6), 1054 (12), 0953 (11), 0217 (3), 1086 (13), 1139 (14), 0208 (2), 0043 (1), 0812 (8), 0587 (7), 0410 (5), 0926 (9), 0948 (10), and 0342 (4). For the Aβ + CU cohort, the corresponding IDs are 0434 (8), 0680 (7), 0075 (13), 0418 (6), 0026 (12), 0206 (5), 0775 (4), 0423 (10), 0841 (2), 0149 (9), 1301 (11), 0768 (3), 0438 (1), and 0296 (14).

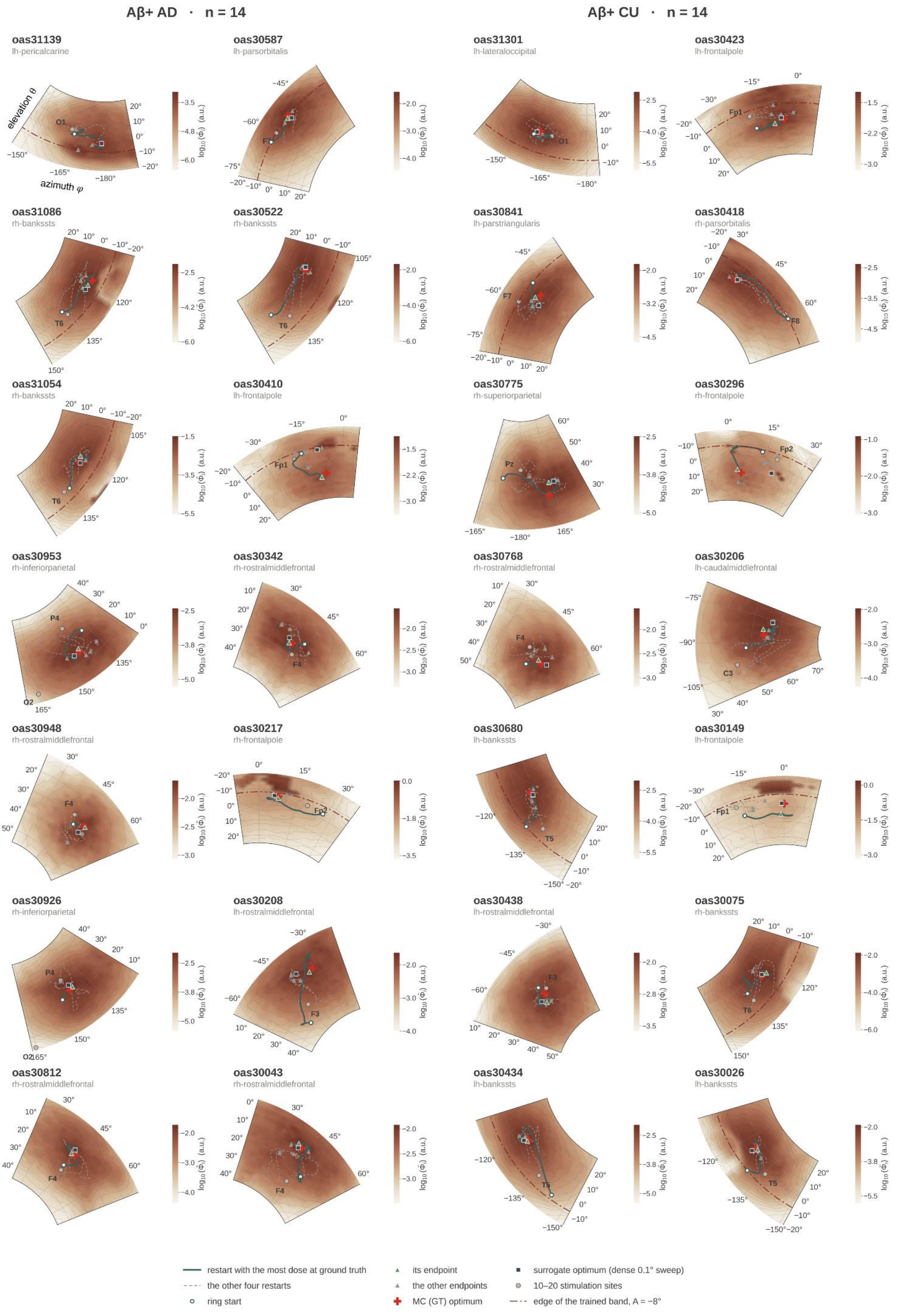


**Fig. S7: Individual optimization paths (n = 28).** Optimization trajectories across subject-specific search sectors, mapped onto a polar projection of MNI space defining the spherical coordinate framework for light source localization. Background: delivered dose (per-panel normalized). Lines: five restarts per participant, with the global optimum as solid and the remaining four as dashed (from initialization to endpoint). Markers: Monte Carlo optimum, dense-sweep surrogate optimum, standard 10–20 sites, and training angular limits. Cohort: amyloid-positive subset with aligned PET-T1w images and Centiloid $\geqslant 26$.

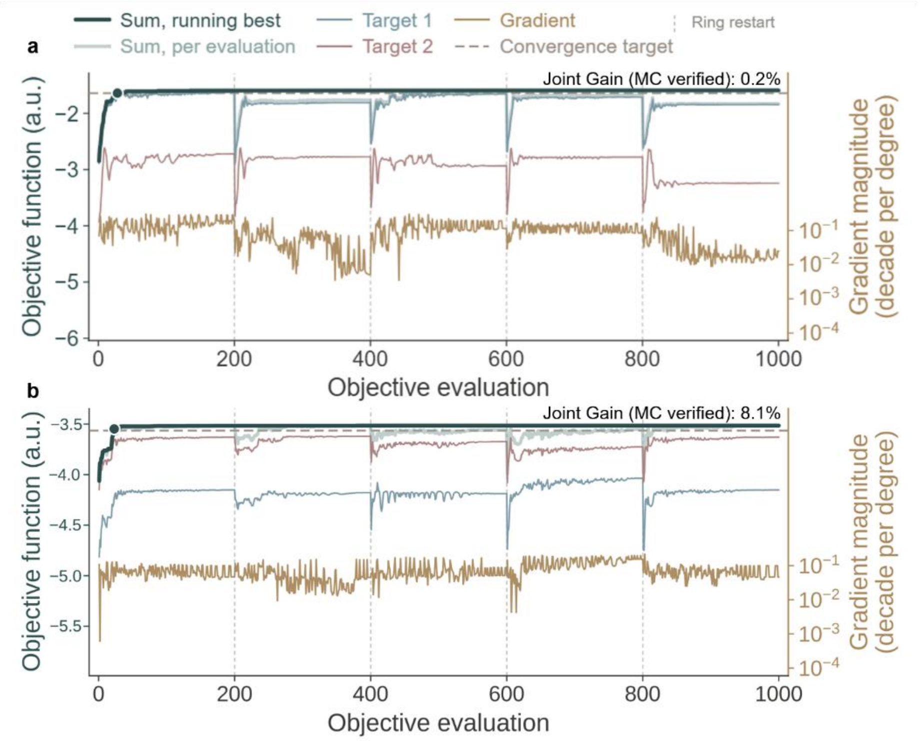


**Fig. S8: Two-target illumination optimization traces.** (**a**) One Aβ+ AD case (oas30812). Single-source Adam optimization against dual targets (1000 calls, six-start ring, linear aggregation). Curves: running objective and its target-specific components; secondary axis: gradient magnitude; rules: dense reference and convergence tolerance. (**b**) One Aβ+ CU case (oas30841). Setup as in (a). Dashed lines: restart boundaries (every 200 calls), where the active restart yields the best surrogate-predicted iterate. In both cases, the voxel distance between the dual targets' local maxima falls below the original 8-mm PET physical resolution, resulting in similar SUVR values and justifying an approximately equal target weighting $(w \approx 1)$ in the objective function.

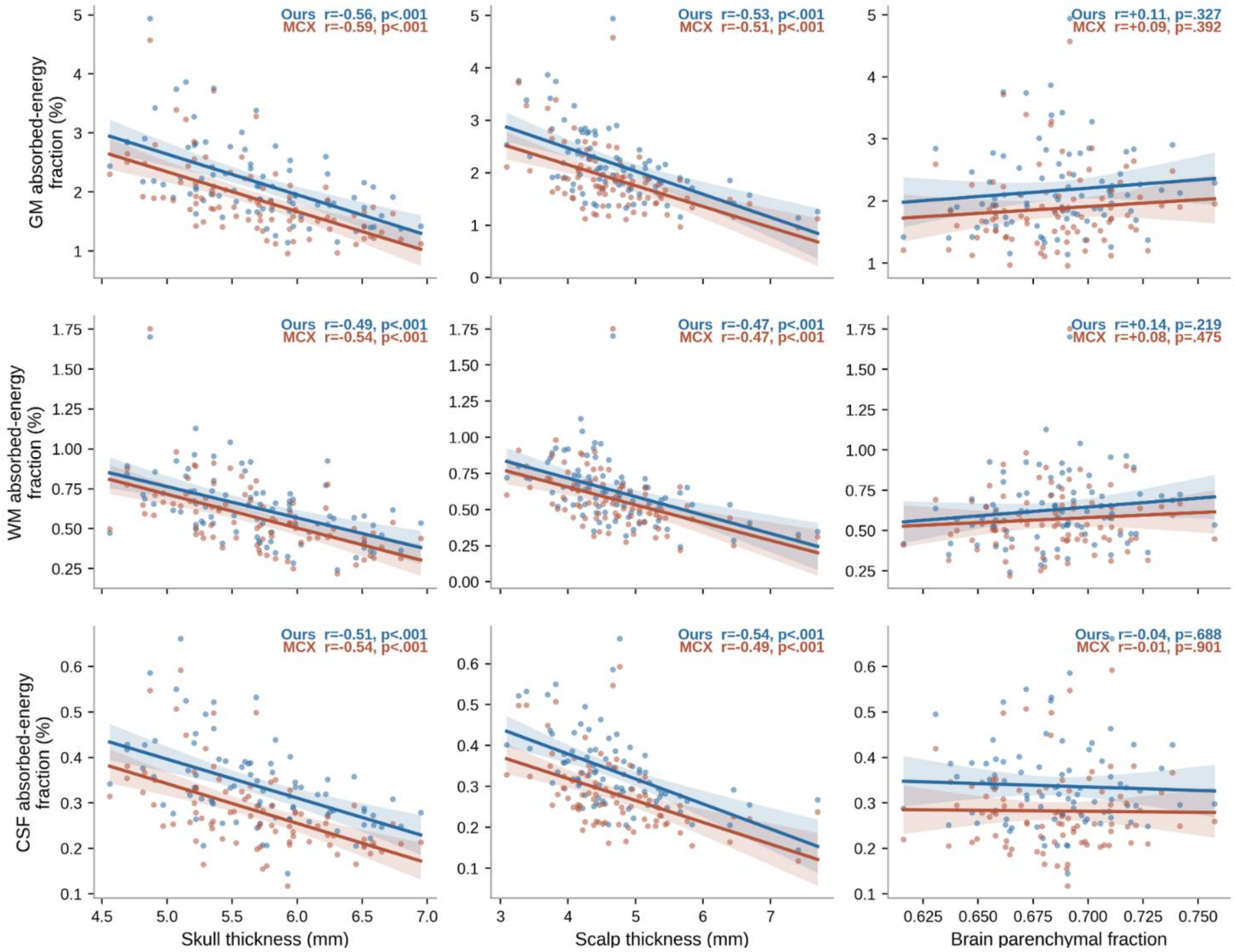


**Fig. S9: Absorbed-energy fraction versus participants' head anatomy (n = 84).** GM, WM and CSF absorbed-energy fractions are plotted versus skull thickness, scalp thickness and brain parenchymal fraction. Blue: PhomiNeuro prediction; red: MC ground truth (calculated via MCX). Values represent per-participant means averaged across the entire standard 10–20 stimulation sites. Scalp thickness is estimated as the sum of skin and muscle. Lines: ordinary least-squares fits; bands: 95% confidence interval of the mean; annotations: Pearson *r*.

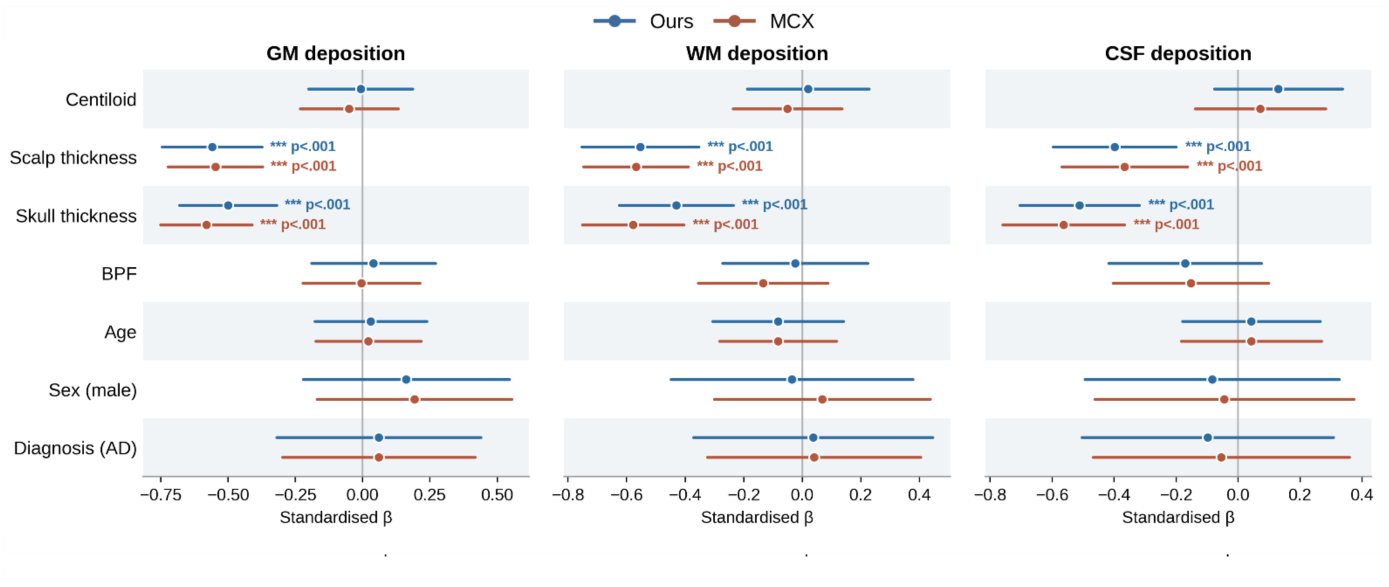


**Fig. S10: : Mutually adjusted predictors of absorbed energy (n = 72).** Standardized regression coefficients (β) from a seven-covariate multivariable model of GM, WM, and CSF absorbed-energy fractions, restricted to participants with Centiloid data. Blue: PhomiNeuro; red: MC. Predictors are z-scored; $\beta$ denotes standard deviations (s.d.) of energy deposition per s.d. of predictor (group differences for sex and diagnosis). Error bars: 95% confidence interval. Scalp and skull thicknesses alone reach statistical significance, with consistent agreement between both methods across all tissue types.

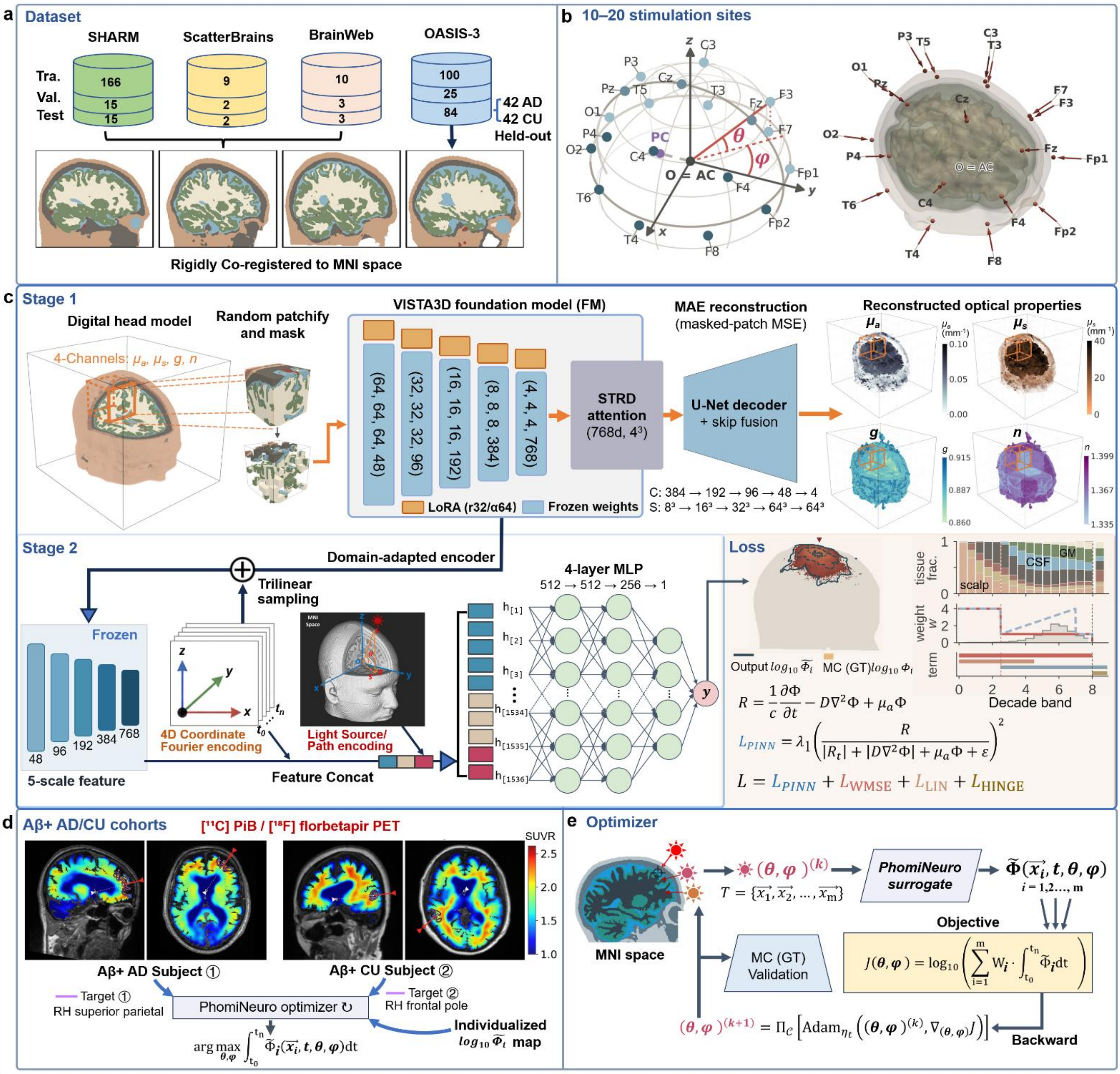


**Figure S11. Details of our proposed PhomiNeuro framework for photon transport surrogate and personalized neuromodulation optimization.** (**a**) Dataset organization and preprocessing. Digital head phantoms from 434 different subjects were assembled from four datasets (SHARM, ScatterBrains, BrainWeb and OASIS-3). Tissue labels were used as provided for the first three sources and derived in this work for OASIS-3 (see Methods). All head phantoms were then registered by a rigid-body transform to a 1 mm isotropic MNI space. Subjects were partitioned without overlap into 330 for training and validation, 20 for development (Dev-test), and 84 for held-out testing. including 42 AD patients and 42 matched cognitively unimpaired (CU) controls, form the held-out test set, and 20 healthy subjects from the other three datasets form the Development (Dev)-test set. The held-out test set includes 42 AD patients and 42 matched cognitively unimpaired (CU) controls. (**b**) Source placement on the international 10–20 system in MNI space. The anterior commissure (AC) is the coordinate origin and the axes follow the MNI frame; source parameters are normalized across subjects and encoded in spherical coordinates ($A$, elevation; $B$, azimuth) derived from the outward scalp normal. (**c**) The architecture design and training of PhomiNeuro: (**i**) Stage1: Domain adaptation of a pretrained 3D medical foundation model (FM) to tissue optics. The encoder is adapted by masked autoencoding with low-rank adaptation (LoRA; rank 32, $\alpha = 64$) on 3D head phantoms with four optical-property channels ($\mu_a$, absorption coefficient; $\mu_s$, scattering coefficient; g, anisotropy factor; n, refractive index) (**ii**) Stage 2:Input encoding and model architecture of the implicit neural representation (INR). For a query coordinate $(x, y, z)$ and time gate $t$, the model predicts $\log_{10} \hat{\Phi}(\boldsymbol{x}, \boldsymbol{y}, \boldsymbol{z}, \boldsymbol{t})$. Input encoding comprises spatiotemporal coordinate features and illumination parameters, including five-scale anatomical features trilinear-sampled from the frozen FM-encoder, band-limited Fourier encodings of $(x, y, z, t)$, local optical properties; and beam direction, beam entry-point coordinate and source-to-point path geometric and optical features, are concatenated and fed to a light-weighted four-layer MLP. (**iii**) Loss function design: The INR loss comprises a physics-informed term $L_{PINN}$ and three data terms. The $L_{PINN}$ is the residual of the time-dependent diffusion equation (TDED), evaluated on seven-point finite-difference collocation stencils. The data terms are a decade-stratified weighted mean-squared error $L_{WMSE}$ against the Monte Carlo (MC) ground truth (GT) in $\log_{10}$ space, a linear-space term $L_{LIN}$ targeting relative error in near-peak region of the light fluence, and a one-sided hinge $L_{HINGE}$ over the MC censored region that penalizes only predictions above the censoring floor. (**d**) Individualized pathology-aware optimization of transcranial photobiomodulation (tPBM) based on amyloid-PET SUVR (n=28, a subset of Aβ+ AD and CU subjects with aligned PET-T1w images and a Centiloid $\geq$ 26). (**e**) Optimizer design for the differentiable PhomINR-based illumination inverse optimization. Abbreviations: MLP, Multi-Layer Perceptron; MNI, Montreal Neurological Institute; AD, Alzheimer's Disease; PET, Positron Emission Tomography; STRD, spatial redundancy dropout; SUVR, Standardized Uptake Value Ratio; T1w, T1-weighted. The 3D phantoms shown are derived from a publicly distributed BrainWeb digital head model and the MNI-ICBM152 average template; neither is a surface reconstruction of an identifiable participant. This figure was created with Python (PyVista, VTK, scikit-image, Matplotlib) and PowerPoint.

**Table S1: Accuracy on the held-out cohort (84 OASIS head models × 6 sites = 504 scenes, gate-averaged, 50 000 sampled points).**

| Model | $R^2$ ↑ | RMSE ↓ | MedAE ↓ | Spearman ρ ↑ | γ (3%/2 px) ↑ | $\lvert\log_{10}$ energy ratio$\rvert$ ↓ |
|---|---|---|---|---|---|---|
| Ours | 0.7385 ± 0.0019 | 0.6211 ± 0.0027 | 0.2466 ± 0.0010 | 0.8923 ± 0.0002 | 0.5899 ± 0.0022 | 0.0281 ± 0.0023 |
| ViT | 0.6973 ± 0.0043 | 0.6934 ± 0.0064 | 0.3059 ± 0.0082 | 0.8901 ± 0.0023 | 0.4665 ± 0.0085 | 0.0257 ± 0.0006 |
| DynUNet | 0.7216 ± 0.0122 | 0.6476 ± 0.0046 | 0.2383 ± 0.0056 | 0.9031 ± 0.0012 | 0.6562 ± 0.0020 | 0.0289 ± 0.0019 |
| SegResNet | 0.7182 ± 0.0079 | 0.6490 ± 0.0037 | 0.2379 ± 0.0047 | 0.9036 ± 0.0012 | 0.6429 ± 0.0100 | 0.0298 ± 0.0061 |
| UNet | 0.7281 ± 0.0026 | 0.6509 ± 0.0009 | 0.2477 ± 0.0015 | 0.9083 ± 0.0015 | 0.5925 ± 0.0033 | 0.0220 ± 0.0018 |
| FNO | 0.5896 ± 0.0185 | 0.7441 ± 0.0061 | 0.3576 ± 0.0072 | 0.8702 ± 0.0030 | 0.3534 ± 0.0074 | 0.0355 ± 0.0024 |
| Coord-RFF | 0.6480 ± 0.0012 | 0.7108 ± 0.0006 | 0.3679 ± 0.0007 | 0.8571 ± 0.0011 | 0.4528 ± 0.0038 | 0.0569 ± 0.0063 |
| SIREN | 0.6291 ± 0.0023 | 0.7320 ± 0.0030 | 0.4012 ± 0.0069 | 0.8548 ± 0.0021 | 0.4140 ± 0.0021 | 0.0658 ± 0.0020 |

Mean ± s.d. over three training seeds; ↑ means higher is better, ↓ means lower is better.

**Table S2: Held-out $R^2$ by time gate.**

| Model | t=0.1 ns | t=0.3 ns | t=0.5 ns | t=0.7 ns | t=0.9 ns | t=1.1 ns | t=1.3 ns | t=1.5 ns | t=1.7 ns | t=1.9 ns | worst gate |
|---|---|---|---|---|---|---|---|---|---|---|---|
| Ours | 0.9242 ± 0.0039 | 0.9383 ± 0.0001 | 0.8462 ± 0.0017 | 0.7409 ± 0.0048 | 0.6693 ± 0.0034 | 0.6507 ± 0.0010 | 0.6623 ± 0.0019 | 0.6678 ± 0.0032 | 0.6419 ± 0.0064 | 0.6438 ± 0.0023 | 0.6410 ± 0.0050 |
| ViT | 0.7191 ± 0.0404 | 0.8834 ± 0.0120 | 0.7429 ± 0.0110 | 0.6246 ± 0.0084 | 0.5992 ± 0.0057 | 0.6288 ± 0.0024 | 0.6719 ± 0.0016 | 0.7010 ± 0.0029 | 0.7070 ± 0.0031 | 0.6948 ± 0.0050 | 0.5992 ± 0.0057 |
| DynUNet | 0.4222 ± 0.1221 | 0.9341 ± 0.0006 | 0.7779 ± 0.0027 | 0.6625 ± 0.0002 | 0.6317 ± 0.0003 | 0.6683 ± 0.0002 | 0.7293 ± 0.0002 | 0.7764 ± 0.0003 | 0.8024 ± 0.0001 | 0.8113 ± 0.0007 | 0.4222 ± 0.1221 |
| SegResNet | 0.3847 ± 0.0685 | 0.9319 ± 0.0014 | 0.7807 ± 0.0042 | 0.6629 ± 0.0026 | 0.6315 ± 0.0009 | 0.6679 ± 0.0019 | 0.7296 ± 0.0012 | 0.7771 ± 0.0015 | 0.8032 ± 0.0011 | 0.8126 ± 0.0016 | 0.3847 ± 0.0685 |
| UNet | 0.5680 ± 0.0319 | 0.9145 ± 0.0023 | 0.7804 ± 0.0011 | 0.6572 ± 0.0007 | 0.6253 ± 0.0009 | 0.6608 ± 0.0005 | 0.7202 ± 0.0001 | 0.7659 ± 0.0004 | 0.7902 ± 0.0012 | 0.7987 ± 0.0021 | 0.5680 ± 0.0319 |
| FNO | -0.1634 ± 0.1858 | 0.8721 ± 0.0023 | 0.7182 ± 0.0034 | 0.6088 ± 0.0011 | 0.5860 ± 0.0010 | 0.6157 ± 0.0020 | 0.6569 ± 0.0037 | 0.6776 ± 0.0048 | 0.6741 ± 0.0048 | 0.6502 ± 0.0028 | -0.1634 ± 0.1858 |
| Coord-RFF | 0.9018 ± 0.0053 | 0.9104 ± 0.0014 | 0.7888 ± 0.0006 | 0.6783 ± 0.0003 | 0.6212 ± 0.0004 | 0.5828 ± 0.0011 | 0.5507 ± 0.0029 | 0.5163 ± 0.0039 | 0.4796 ± 0.0038 | 0.4499 ± 0.0036 | 0.4499 ± 0.0036 |
| SIREN | 0.9045 ± 0.0160 | 0.8829 ± 0.0085 | 0.7747 ± 0.0024 | 0.6650 ± 0.0018 | 0.6091 ± 0.0003 | 0.5653 ± 0.0024 | 0.5265 ± 0.0029 | 0.4912 ± 0.0028 | 0.4556 ± 0.0012 | 0.4159 ± 0.0054 | 0.4159 ± 0.0054 |

Mean ± across-seed s.d. over three training seeds. t is the gate center in ns. The worst-gate column is each seed's own weakest gate.

**Table S3: Held-out RMSE gate by gate.**

| Model | t=0.1 | t=0.3 | t=0.5 | t=0.7 | t=0.9 | t=1.1 | t=1.3 | t=1.5 | t=1.7 | t=1.9 |
|---|---|---|---|---|---|---|---|---|---|---|
| Ours | 0.1350 ± 0.0023 | 0.3406 ± 0.0002 | 0.7704 ± 0.0044 | 1.0755 ± 0.0105 | 1.0585 ± 0.0059 | 0.8326 ± 0.0011 | 0.6244 ± 0.0015 | 0.5055 ± 0.0021 | 0.4555 ± 0.0032 | 0.4132 ± 0.0009 |
| ViT | 0.2614 ± 0.0212 | 0.4622 ± 0.0220 | 0.9931 ± 0.0212 | 1.2966 ± 0.0145 | 1.1686 ± 0.0080 | 0.8619 ± 0.0029 | 0.6175 ± 0.0011 | 0.4807 ± 0.0015 | 0.4123 ± 0.0012 | 0.3798 ± 0.0013 |
| DynUNet | 0.3930 ± 0.0441 | 0.3529 ± 0.0014 | 0.9268 ± 0.0057 | 1.2314 ± 0.0004 | 1.1212 ± 0.0005 | 0.8172 ± 0.0003 | 0.5646 ± 0.0003 | 0.4200 ± 0.0002 | 0.3441 ± 0.0001 | 0.3048 ± 0.0003 |
| SegResNet | 0.4076 ± 0.0198 | 0.3590 ± 0.0039 | 0.9216 ± 0.0088 | 1.2310 ± 0.0048 | 1.1218 ± 0.0015 | 0.8178 ± 0.0023 | 0.5643 ± 0.0013 | 0.4195 ± 0.0013 | 0.3436 ± 0.0009 | 0.3041 ± 0.0011 |
| UNet | 0.3176 ± 0.0148 | 0.3996 ± 0.0041 | 0.9217 ± 0.0024 | 1.2412 ± 0.0012 | 1.1310 ± 0.0014 | 0.8262 ± 0.0006 | 0.5736 ± 0.0001 | 0.4294 ± 0.0002 | 0.3542 ± 0.0007 | 0.3144 ± 0.0012 |
| FNO | 0.5505 ± 0.0545 | 0.4871 ± 0.0049 | 1.0426 ± 0.0065 | 1.3250 ± 0.0019 | 1.1883 ± 0.0016 | 0.8771 ± 0.0023 | 0.6314 ± 0.0034 | 0.4987 ± 0.0040 | 0.4345 ± 0.0036 | 0.4054 ± 0.0025 |
| Coord-RFF | 0.1580 ± 0.0037 | 0.4105 ± 0.0030 | 0.9046 ± 0.0016 | 1.1999 ± 0.0004 | 1.1323 ± 0.0007 | 0.9064 ± 0.0009 | 0.7174 ± 0.0020 | 0.6100 ± 0.0021 | 0.5524 ± 0.0017 | 0.5167 ± 0.0014 |
| SIREN | 0.1577 ± 0.0107 | 0.4665 ± 0.0165 | 0.9351 ± 0.0048 | 1.2255 ± 0.0032 | 1.1515 ± 0.0006 | 0.9253 ± 0.0025 | 0.7362 ± 0.0025 | 0.6257 ± 0.0017 | 0.5646 ± 0.0008 | 0.5323 ± 0.0020 |

Mean ± across-seed s.d. over three training seeds. t is the time gate center in ns.

**Table S4: Identical checkpoints and evaluation protocol on the development and the held-out cohort.**

| Model | $R^2$ (Dev) | RMSE (Dev) | medAE (Dev) | $R^2$ (held-out) | RMSE (held-out) | medAE (held-out) | $\Delta R^2$ |
|---|---|---|---|---|---|---|---|
| Ours | 0.7721 ± 0.0008 | 0.5043 ± 0.0019 | 0.2200 ± 0.0015 | 0.7385 ± 0.0019 | 0.6211 ± 0.0027 | 0.2466 ± 0.0010 | -0.0336 |
| ViT | 0.7263 ± 0.0033 | 0.5797 ± 0.0038 | 0.2914 ± 0.0032 | 0.6973 ± 0.0043 | 0.6934 ± 0.0064 | 0.3059 ± 0.0082 | -0.0290 |
| DynUNet | 0.8183 ± 0.0048 | 0.4908 ± 0.0031 | 0.1975 ± 0.0043 | 0.7216 ± 0.0122 | 0.6476 ± 0.0046 | 0.2383 ± 0.0056 | -0.0967 |
| SegResNet | 0.8216 ± 0.0012 | 0.4892 ± 0.0004 | 0.1972 ± 0.0037 | 0.7182 ± 0.0079 | 0.6490 ± 0.0037 | 0.2379 ± 0.0047 | -0.1033 |
| UNet | 0.8211 ± 0.0016 | 0.4940 ± 0.0019 | 0.2037 ± 0.0021 | 0.7281 ± 0.0026 | 0.6509 ± 0.0009 | 0.2477 ± 0.0015 | -0.0930 |
| FNO | 0.6988 ± 0.0031 | 0.5993 ± 0.0021 | 0.3179 ± 0.0041 | 0.5896 ± 0.0185 | 0.7441 ± 0.0061 | 0.3576 ± 0.0072 | -0.1092 |
| Coord-RFF | 0.6021 ± 0.0023 | 0.6489 ± 0.0009 | 0.3259 ± 0.0009 | 0.6480 ± 0.0012 | 0.7108 ± 0.0006 | 0.3679 ± 0.0007 | +0.0459 |
| SIREN | 0.5937 ± 0.0053 | 0.6600 ± 0.0024 | 0.3499 ± 0.0034 | 0.6291 ± 0.0023 | 0.7320 ± 0.0030 | 0.4012 ± 0.0069 | +0.0354 |

Mean ± across-seed s.d., three seeds. Same checkpoints, no retraining; 50 000 query coordinatess per scene. Development (Dev), 20 heads × 19 sites = 380 scenes; held-out, 84 heads × 6 electrodes (a subset of the 10-20 stimulation site) = 504 scenes. $\Delta R^2$ mixes a cohort effect with an electrode effect.

### Table S5: Ablation of the developed model on the held-out cohort

| Variant | $R^2$ ↑ | RMSE ↓ | MedAE ↓ | Spearman ρ ↑ | DICE ↑ | CCC ↑ | γ (3%/2 px) ↑ |
|---|---|---|---|---|---|---|---|
| Ours | 0.7109 ± 0.0013 | 0.6495 ± 0.0016 | 0.2872 ± 0.0028 | 0.8827 ± 0.0012 | 0.8853 ± 0.0009 | 0.8455 ± 0.0006 | 0.5601 ± 0.0017 |
| − FM encoder | 0.6435 ± 0.0016 | 0.6962 ± 0.0014 | 0.2997 ± 0.0029 | 0.8578 ± 0.0013 | 0.8992 ± 0.0011 | 0.8200 ± 0.0008 | 0.4063 ± 0.0063 |
| FM encoder (not fine-tuned) | 0.6989 ± 0.0018 | 0.6554 ± 0.0038 | 0.2936 ± 0.0156 | 0.8822 ± 0.0048 | 0.8883 ± 0.0085 | 0.8417 ± 0.0010 | 0.5122 ± 0.0043 |
| Random encoder | 0.6250 ± 0.0080 | 0.7007 ± 0.0122 | 0.3247 ± 0.0317 | 0.8702 ± 0.0091 | 0.8970 ± 0.0164 | 0.8216 ± 0.0038 | 0.4846 ± 0.0097 |
| − path integrals | 0.6520 ± 0.0138 | 0.6821 ± 0.0093 | 0.3045 ± 0.0132 | 0.8822 ± 0.0021 | 0.8972 ± 0.0025 | 0.8256 ± 0.0050 | 0.4961 ± 0.0026 |
| − loss term $L_{PINN}$ | 0.7176 ± 0.0018 | 0.6491 ± 0.0001 | 0.3031 ± 0.0014 | 0.8888 ± 0.0005 | 0.8885 ± 0.0013 | 0.8453 ± 0.0001 | 0.5755 ± 0.0014 |
| − loss term $L_{lin}$ | 0.7138 ± 0.0005 | 0.6461 ± 0.0012 | 0.2840 ± 0.0033 | 0.8883 ± 0.0012 | 0.8902 ± 0.0037 | 0.8459 ± 0.0007 | 0.5509 ± 0.0026 |
| − loss term $L_{hinge}$ | 0.2375 ± 0.0295 | 1.1157 ± 0.0223 | 0.7065 ± 0.0357 | 0.6986 ± 0.0074 | 0.7575 ± 0.0000 | 0.5898 ± 0.0095 | 0.2072 ± 0.0833 |
| − PINN and − FM encoder | 0.6618 ± 0.0016 | 0.6987 ± 0.0017 | 0.3421 ± 0.0026 | 0.8582 ± 0.0007 | 0.8850 ± 0.0007 | 0.8159 ± 0.0008 | 0.4177 ± 0.0050 |

Mean ± across-seed s.d. over three training seeds. A difference from the complete model is outside the noise when it exceeds $1.96\sqrt{((\sigma_h^2 + \sigma_v^2)/3)}$.

### Table S6: Within-group comparison of male v.s female participants of the OASIS-3 test cohort.

| Characteristic | Male | Female | P |
|---|---|---|---|
| **AD (male n = 23, female n = 19)** | | | |
| Age (years) | 78.5 ± 7.6 | 75.2 ± 7.4 | 0.093 |
| Education (years) | 16.7 ± 2.8 | 14.1 ± 2.5 | **0.007** |
| MMSE (/30) | 26.5 ± 3.1 | 26.2 ± 3.3 | 0.759 |
| CDR-SB (/18) [b] | 4.0 ± 2.6 | 2.8 ± 2.4 | 0.094 |
| Centiloid [b] | 53.0 ± 44.3 | 62.5 ± 43.2 | 0.436 |
| Scalp thickness, skin + muscle (mm) | 5.03 ± 1.00 | 4.16 ± 0.50 | **0.001** |
| Skull thickness (mm) | 5.76 ± 0.50 | 5.61 ± 0.64 | 0.277 |
| BPF | 0.671 ± 0.023 | 0.684 ± 0.030 | 0.300 |
| GM absorbed-energy fraction, surrogate (%) | 2.181 ± 0.814 | 2.337 ± 0.699 | 0.245 |
| GM absorbed-energy fraction, Monte Carlo (%) | 1.886 ± 0.790 | 2.036 ± 0.652 | 0.206 |
| WM absorbed-energy fraction, surrogate (%) | 0.638 ± 0.286 | 0.699 ± 0.206 | 0.157 |
| WM absorbed-energy fraction, Monte Carlo (%) | 0.586 ± 0.301 | 0.633 ± 0.189 | 0.180 |
| CSF absorbed-energy fraction, surrogate (%) | 0.338 ± 0.098 | 0.355 ± 0.079 | 0.266 |
| CSF absorbed-energy fraction, Monte Carlo (%) | 0.283 ± 0.096 | 0.299 ± 0.075 | 0.216 |
| APOE ε4 carrier | 11/23 (48%) | 12/19 (63%) | 0.366 |
| **CU (male n = 23, female n = 19)** | | | |
| Age (years) | 78.6 ± 7.8 | 75.3 ± 7.5 | 0.095 |
| Education (years) | 16.4 ± 2.0 | 15.5 ± 2.5 | 0.247 |
| MMSE (/30) | 28.6 ± 1.5 | 28.7 ± 1.1 | 0.927 |
| CDR-SB (/18) | 0.0 ± 0.1 | 0.0 ± 0.1 | 0.918 |
| Centiloid | 22.0 ± 27.6 | 27.3 ± 30.2 | 0.505 |

| Characteristic | Male | Female | P |
|---|---|---|---|
| Scalp thickness, skin + muscle (mm) | 4.96 ± 0.74 | 4.48 ± 0.61 | **0.042** |
| Skull thickness (mm) | 5.75 ± 0.52 | 5.61 ± 0.58 | 0.306 |
| BPF | 0.684 ± 0.021 | 0.704 ± 0.026 | **0.008** |
| GM absorbed-energy fraction, surrogate (%) | 1.930 ± 0.617 | 2.264 ± 0.493 | **0.002** |
| GM absorbed-energy fraction, Monte Carlo (%) | 1.683 ± 0.568 | 1.942 ± 0.434 | **0.012** |
| WM absorbed-energy fraction, surrogate (%) | 0.536 ± 0.163 | 0.660 ± 0.180 | **0.028** |
| WM absorbed-energy fraction, Monte Carlo (%) | 0.496 ± 0.167 | 0.579 ± 0.151 | 0.086 |
| CSF absorbed-energy fraction, surrogate (%) | 0.300 ± 0.089 | 0.363 ± 0.097 | **0.016** |
| CSF absorbed-energy fraction, Monte Carlo (%) | 0.250 ± 0.085 | 0.303 ± 0.092 | **0.038** |
| APOE ε4 carrier | 3/23 (13%) | 4/19 (21%) | 0.682 |
| **Aβ-positive (CL ≥ 26) (male n = 17, female n = 17)** | | | |
| Age (years) | 82.6 ± 6.0 | 77.7 ± 6.2 | **0.006** |
| Education (years) | 16.8 ± 2.5 | 15.1 ± 2.7 | 0.085 |
| MMSE (/30) | 27.1 ± 2.9 | 27.1 ± 3.0 | 1.000 |
| CDR-SB (/18) [b] | 2.9 ± 2.9 | 1.9 ± 2.8 | 0.292 |
| Centiloid | 76.4 ± 23.6 | 73.4 ± 29.8 | 0.809 |
| Scalp thickness, skin + muscle (mm) | 4.77 ± 0.61 | 4.14 ± 0.49 | **0.004** |
| Skull thickness (mm) | 5.74 ± 0.52 | 5.61 ± 0.52 | 0.581 |
| BPF | 0.667 ± 0.024 | 0.690 ± 0.031 | **0.036** |
| GM absorbed-energy fraction, surrogate (%) | 2.157 ± 0.559 | 2.245 ± 0.486 | 0.535 |
| GM absorbed-energy fraction, Monte Carlo (%) | 1.842 ± 0.504 | 1.925 ± 0.385 | 0.449 |
| WM absorbed-energy fraction, surrogate (%) | 0.608 ± 0.171 | 0.685 ± 0.166 | 0.242 |
| WM absorbed-energy fraction, Monte Carlo (%) | 0.547 ± 0.152 | 0.604 ± 0.124 | 0.335 |
| CSF absorbed-energy fraction, surrogate (%) | 0.345 ± 0.064 | 0.377 ± 0.091 | 0.256 |
| CSF absorbed-energy fraction, Monte Carlo (%) | 0.282 ± 0.058 | 0.314 ± 0.085 | 0.256 |
| APOE ε4 carrier | 10/17 (59%) | 10/17 (59%) | 1.000 |
| **Aβ-negative (CL < 26) (male n = 23, female n = 15)** | | | |
| Age (years) | 76.0 ± 8.2 | 72.7 ± 7.0 | 0.179 |
| Education (years) | 16.4 ± 2.3 | 14.6 ± 2.7 | **0.038** |
| MMSE (/30) | 28.4 ± 1.8 | 28.5 ± 1.6 | 0.914 |
| CDR-SB (/18) [b] | 1.0 ± 2.0 | 0.6 ± 1.2 | 0.972 |
| Centiloid | 6.1 ± 8.5 | 7.9 ± 8.9 | 0.446 |
| Scalp thickness, skin + muscle (mm) | 5.09 ± 1.02 | 4.65 ± 0.58 | 0.199 |
| Skull thickness (mm) | 5.83 ± 0.51 | 5.66 ± 0.73 | 0.420 |
| BPF | 0.682 ± 0.022 | 0.704 ± 0.026 | **0.012** |
| GM absorbed-energy fraction, surrogate (%) | 1.882 ± 0.643 | 2.218 ± 0.617 | **0.020** |
| GM absorbed-energy fraction, Monte Carlo (%) | 1.662 ± 0.625 | 1.917 ± 0.555 | 0.073 |
| WM absorbed-energy fraction, surrogate (%) | 0.514 ± 0.168 | 0.624 ± 0.173 | 0.056 |
| WM absorbed-energy fraction, Monte Carlo (%) | 0.484 ± 0.183 | 0.555 ± 0.169 | 0.170 |
| CSF absorbed-energy fraction, surrogate (%) | 0.302 ± 0.099 | 0.330 ± 0.077 | 0.094 |
| CSF absorbed-energy fraction, Monte Carlo (%) | 0.257 ± 0.098 | 0.276 ± 0.074 | 0.221 |
| APOE ε4 carrier | 2/23 (9%) | 3/15 (20%) | 0.365 |

Data are mean ± s.d. or n (%). Within each stratum, males and females were compared with the Mann–Whitney U test (continuous variables) or Fisher's exact test (APOE ε4); Amyloid strata comprise the participants with an amyloid-PET scan (Aβ+ n = 34, Aβ− n = 38 of 72 scanned); the AD and CU strata use all 42 participants each, except for Centiloid. Thickness values are means over that participant's 10–20 stimulation sites; absorbed-energy fractions are means over the 10–20 sites.

**Table S7: Within-diagnosis comparison of amyloid-positive v.s amyloid-negative participants of the OASIS-3 test cohort.**

| Characteristic | Aβ-positive | Aβ-negative | P |
|---|---|---|---|
| **AD (Aβ+ n = 20, Aβ− n = 12)** | | | |
| Sex (male / female) | 11 / 9 | 7 / 5 | 1.000 |
| Age (years) | 79.3 ± 6.6 | 73.7 ± 8.2 | 0.070 |
| Education (years) | 15.5 ± 2.9 | 15.7 ± 3.2 | 0.829 |
| MMSE (/30) | 25.9 ± 3.3 | 28.1 ± 2.1 | 0.072 |
| CDR-SB (/18) | 4.1 ± 2.6 | 2.5 ± 2.3 | 0.051 |
| Centiloid | 87.3 ± 21.8 | 7.0 ± 8.9 | **<0.001** |
| Scalp thickness, skin + muscle (mm) | 4.42 ± 0.68 | 5.03 ± 1.19 | 0.070 |
| Skull thickness (mm) | 5.77 ± 0.60 | 5.69 ± 0.59 | 0.861 |
| BPF | 0.670 ± 0.029 | 0.685 ± 0.027 | 0.167 |
| GM absorbed-energy fraction, surrogate (%) | 2.181 ± 0.588 | 2.022 ± 0.631 | 0.340 |
| GM absorbed-energy fraction, Monte Carlo (%) | 1.863 ± 0.500 | 1.781 ± 0.638 | 0.403 |
| WM absorbed-energy fraction, surrogate (%) | 0.630 ± 0.161 | 0.575 ± 0.194 | 0.546 |

| Characteristic | Aβ-positive | Aβ-negative | P |
|---|---|---|---|
| WM absorbed-energy fraction, Monte Carlo (%) | 0.563 ± 0.136 | 0.531 ± 0.204 | 0.599 |
| CSF absorbed-energy fraction, surrogate (%) | 0.357 ± 0.062 | 0.316 ± 0.093 | 0.064 |
| CSF absorbed-energy fraction, Monte Carlo (%) | 0.293 ± 0.056 | 0.271 ± 0.094 | 0.167 |
| APOE ε4 carrier | 15/20 (75%) | 3/12 (25%) | **0.010** |
| **CU (Aβ+ n = 14, Aβ− n = 26)** | | | |
| Sex (male / female) | 6 / 8 | 16 / 10 | 0.327 |
| Age (years) | 81.2 ± 6.3 | 75.2 ± 7.8 | **0.023** |
| Education (years) | 16.5 ± 2.3 | 15.7 ± 2.3 | 0.356 |
| MMSE (/30) | 28.7 ± 1.1 | 28.6 ± 1.5 | 0.965 |
| CDR-SB (/18) | 0.0 ± 0.0 | 0.0 ± 0.1 | 0.311 |
| Centiloid | 57.2 ± 22.8 | 6.7 ± 8.6 | **<0.001** |
| Scalp thickness, skin + muscle (mm) | 4.50 ± 0.58 | 4.87 ± 0.74 | 0.137 |
| Skull thickness (mm) | 5.55 ± 0.35 | 5.79 ± 0.62 | 0.109 |
| BPF | 0.691 ± 0.027 | 0.693 ± 0.025 | 0.590 |
| GM absorbed-energy fraction, surrogate (%) | 2.229 ± 0.416 | 2.011 ± 0.666 | 0.059 |
| GM absorbed-energy fraction, Monte Carlo (%) | 1.912 ± 0.364 | 1.754 ± 0.601 | 0.103 |
| WM absorbed-energy fraction, surrogate (%) | 0.671 ± 0.187 | 0.549 ± 0.171 | 0.059 |
| WM absorbed-energy fraction, Monte Carlo (%) | 0.593 ± 0.148 | 0.503 ± 0.170 | 0.056 |
| CSF absorbed-energy fraction, surrogate (%) | 0.366 ± 0.101 | 0.312 ± 0.091 | **0.040** |
| CSF absorbed-energy fraction, Monte Carlo (%) | 0.305 ± 0.095 | 0.261 ± 0.088 | 0.059 |
| APOE ε4 carrier | 5/14 (36%) | 2/26 (8%) | **0.039** |

Data are mean ± s.d. or n (%). Restricted to the 72 participants with an amyloid-PET scan. Amyloid positivity is Centiloid ≥ 26. Within each diagnosis, Aβ+ cohort and Aβ− cohort were compared with the Mann–Whitney U test (continuous variables) or Fisher's exact test (sex, APOE ε4); bold, $P < 0.05$. Absorbed-energy fractions are the share of the head's total absorbed energy deposited in each tissue, averaged over the 10-20 sites. AD, Alzheimer's disease; CU, cognitively unimpaired; BPF, brain parenchymal fraction; GM, grey matter; WM, white matter; CSF, cerebrospinal fluid; CDR-SB, Clinical Dementia Rating–Sum of Boxes; MMSE, Mini-Mental State Examination; APOE, apolipoprotein E; Aβ, amyloid-β; CL, Centiloid.

**Video S1: Time-resolved 3D light fluence propagation of PhomiNeuro prediction versus Monte Carlo ground truth.** Sequential animation across ten 0.2 ns gates (t = 0.1–1.9 ns) displayed at 2 fps. A dev-test head model (scb15) is displayed via concentric, semi-transparent tissue envelopes (scalp, skull, CSF, GM, WM). Fluence is visualized as a particle cloud with amplitude encoded by particle color across eight decades. red arrow: beam direction at Fz.

**Video S2: Continuous-time frame interpolation between supervised time gates.** Temporal interpolation performance at 4 fps (head sh076 under Fz illumination). Top row: Each 0.2 ns gate held for four frames. Left: PhomiNeuro; right: baseline $5\times10^7$ MC. Bottom row: continuous-time interpolation with 0.05 ns steps. Left: PhomiNeuro temporal interpolation; right: $5\times10^9$ MC reference. Light fluence is integrated over a 0.2 ns window; red arrow: beam direction.
